# ABI5 Binding Proteins are substrates of key components in the ABA core signaling pathway

**DOI:** 10.1101/2024.10.11.617944

**Authors:** Tim J Lynch, B. Joy Erickson McNally, Teodora Losic, Jonas Lindquist, Ruth Finkelstein

**Affiliations:** Department of Molecular, Cellular and Developmental Biology, University of California at Santa Barbara, Santa Barbara, CA 93106; Biomolecular Science and Engineering Program, University of California at Santa Barbara, CA 93106

## Abstract

The central components of the ABA core signaling pathway are families of receptors, clade A type 2C protein phosphatases (PP2Cs), SNF1-Related Protein Kinases (SnRK2s), and diverse sets of proteins regulated by phosphorylation via these kinases, including bZIP transcription factors such as ABA-INSENSITIVE(ABI)5. The larger network of ABA signaling factors includes additional kinases and E3 ligases that modify these components to affect their activity and stability. The ABI5-Binding Proteins (AFPs) are negative regulators of ABA response. This study shows that the AFPs interact with specific family members of all components of this pathway and are substrates for SnRK2s and PP2Cs. AFPs also interact with subsets of MAP kinases (MPKs) and 14-3-3 proteins previously found to regulate activity of the ABI5-related clade of transcription factors. Residues predicted to be phosphorylated are conserved between AFPs, but are located within regions predicted to be unstructured. ABA promotes phosphorylation of AFP2, but conditions that prevent phosphorylation of AFP2 result in decreased stability, a shift in localization toward dispersed foci, and reduced effectiveness for inhibiting ABA response at germination. Thus, AFP2 appears to be an important hub in the ABA core signaling pathway.

**One-Sentence Summary:** An inhibitor of ABA-INSENSITIVE5 is proposed to act as a hub in ABA signaling, and its stability, localization and function are regulated by changes in phosphorylation state.

## INTRODUCTION

The phytohormone ABA regulates diverse responses in plants, in part mediated by large changes in gene expression. Although transcriptome experiments in diverse tissues at various stages of growth have identified many thousands of ABA-regulated genes, very few of these are consistently ABA responsive across all of these conditions (reviewed in Finkelstein, 2013) and most transcriptome comparisons focus on co-expressed genes to construct gene regulatory networks acting in specific responses or tissues (reviewed in Ko and Brandizzi, 2020). However, some loci encode core components regulating these diverse responses. The central components of the ABA core signaling pathway are families of receptors, protein phosphatases, protein kinases, and diverse sets of proteins regulated by phosphorylation via these kinases (Cutler et al., 2010; Chen et al., 2020; Ali et al., 2020). In the absence of ABA, PP2C clade A protein phosphatases bind to SNF1-Related Protein Kinase2s (SnRK2s), sterically preventing them from becoming phosphorylated and active. When ABA is present, PYRABACTIN RESISTANT (PYR/PYL)/REGULATORY COMPONENT OF ABA RECEPTOR (RCAR) receptors bind and inactivate these PP2C phosphatases, thereby permitting phosphorylation and activation of the SnRK2 class kinases. These kinases can then phosphorylate regulatory proteins such as the ABA-INSENSITIVE(ABI)5/ABA RESPONSE ELEMENT BINDING FACTOR (AREB/ABF) clade of bZIP transcription factors, leading to ABA-induced changes in gene expression, including feedback regulation by inducing expression of the PP2C co-receptors (Wang et al., 2019). Several of these bZIP proteins are bound in a phosphorylation-dependent manner by 14-3-3 proteins, and this interaction can be required for gene activation by the bZIP factor (Schoonheim et al., 2007; Camoni et al., 2018).

The clade A PP2Cs are monomeric Ser/Thr phosphatases that were initially identified genetically by dominant mutations in ABA-INSENSITIVE(ABI)1 and ABI2 (Koornneef et al., 1984), resulting in ABA resistance due to their inability to be inactivated (Park et al., 2009; reviewed in Finkelstein, 2013). Additional members of the clade, e.g. ABA-HYPERSENSITIVE GERMINATION(AHG)1 and AHG3, were identified based on loss of function mutations conferring hypersensitivity to ABA, reflecting their role as negative regulators of ABA response (Yoshida et al., 2006; Nishimura et al., 2007). These PP2Cs have overlapping expression patterns and functions, such that double mutants are more hypersensitive than the single mutants. Interactome and other studies have identified numerous substrates or other interaction partners for these PP2Cs, including members of multiple families of kinases and transcription factors involved in response to ABA and abiotic stresses (Lumba et al., 2014; Yilmaz et al., 2022).

The SnRK2 Ser/Thr kinases are a plant-specific subfamily of the SnRKs, comprising 10 family members in *Arabidopsis thaliana* (reviewed in Kulik et al., 2011; Maszkowska et al., 2021). Although all SnRK2s function in stress responses, this subfamily is further divided into 3 groups distinguished by their response to ABA. Group 1 does not respond to ABA, group 2 shows minimal response to ABA, and group 3 is strongly activated by ABA (Kobayashi et al., 2004). In Arabidopsis, group 3 consists of SnRK2.2, SnRK2.3 and SnRK2.6. These kinases have partially redundant functions, with SnRK2.2 and SnRK2.3 playing major roles in seed dormancy, control of germination and seedling growth, while SnRK2.6 has greater effects on stomatal regulation (Fujii and Zhu, 2009; Nakashima et al., 2009). Triple mutants defective in all three are highly resistant to ABA and exhibit extreme drought sensitivity and severely stunted growth (Fujita et al., 2009). In contrast, SnRK2.10 is a member of group 1 and responds to osmotic stress, but is not part of the ABA core signaling pathway (Kobayashi et al., 2004).

Mitogen activated protein kinases (MAPKs, also known as MPKs) are another conserved family of Ser/Thr kinases, some of which regulate abiotic stress and ABA response. MAPKs are the final step in the MAPK cascade, composed of MAP kinase kinase kinases (MAPKKKs/MAP3Ks/MEKKs) that phosphorylate and activate MAP kinase kinases (MAPKKs/MAP2Ks/MEKs), which in turn phosphorylate/activate the MAPKs (Jagodzik et al., 2018). Over 100 genes encode members of these families in *Arabidopsis thaliana,* regulating diverse processes with substantial overlaps in function. Although not part of the ABA core signaling pathway, cross-talk between these pathways occurs at various levels: SnRK2s phosphorylate some MAPKs (Umezawa et al., 2013), while MAP3Ks phosphorylate SnRK2s and are required for their activation by ABA (Takahashi et al., 2020). MAPK cascades regulate ABA response through effects on gene expression, including induction of transcription factors such as ABI3 and ABI5 (Lee et al., 2014), and activation or stabilization of other factors, e.g. ABI4, by phosphorylation (Guo et al., 2016; Eisner et al., 2021; Maymon et al., 2022). The 20 MAPKs of Arabidopsis are divided into 4 clades (A-D), distinguished by their domain structures and the sequence at the site phosphorylated by MAPKKs, which is different in clade D (Group et al., 2002). Clade D also lacks a “common docking” domain that is recognized by MAPKKs, phosphatases and other proteins. The most extensively studied MAPKs are MPK3, MPK4 and MPK6, which are members of clades A and B; these have been shown to function in diverse pathways including stomatal development, root growth, pollen tube growth, immune response, and response to abiotic stress and/or ABA (Wang et al., 2007; Shao et al., 2020; Galletti et al., 2011; Zou et al., 2021; Guan et al., 2014; de Zelicourt et al., 2016). All members of the C clade are also activated by ABA by inducing synthesis of the upstream MAP3Ks via the PYR/PYL/RCAR-PP2C-SnRK2 core signaling pathway (Danquah et al., 2015). Within this clade, MPK7 has been specifically implicated in dormancy release (Chen et al., 2023).

Although not initially characterized as part of the core signaling pathway, a group of ABI5-binding proteins (AFPs) identified by a yeast two-hybrid screen using ABI5 as bait were shown to be important ABA-inducible negative regulators of ABA response (Garcia et al., 2008). One of these, AFP3, is among the few genes that are ABA-induced in many different contexts, but especially during vegetative growth (Finkelstein, 2013). In contrast, AFP1 and AFP2 are most active during germination and seedling establishment. The AFPs have three highly conserved domains, including an <u>e</u>thylene-responsive element binding factor-associated <u>a</u>mphiphilic repression (EAR) domain in the “A domain” and a nuclear localization signal in the “B domain,” in addition to the C-terminal domain initially shown to interact with the ABI5/ABF/AREB transcription factors (Garcia et al., 2008). Overexpression of the AFPs results in extreme ABA resistance, including decreased expression of many seed maturation genes, failure to acquire desiccation tolerance, and ability to germinate on 100-fold higher ABA concentrations than wild-type seeds (Lynch et al., 2017).

Studies addressing the mechanism by which AFPs inhibit ABA response have demonstrated interactions with co-repressors including TOPLESS/TOPLESS-RELATED (TPL/TPR) and Histone Deacetylase (HDAC) subunits, suggesting effects on chromatin remodeling (Lynch et al., 2017). Although mechanistically more obscure, the AFPs also interact directly with DELLA proteins, antagonizing their inhibitory effects on germination (Finkelstein and Lynch, 2022). The AFPs have also been proposed to confer ABA resistance by promoting ABI5 degradation, functioning as adaptors for E3 ligases that ubiquitinate ABI5 and possibly related bZIP proteins, based in part on colocalization of AFP1 with ABI5 and COP1 in nuclear bodies (Lopez-Molina et al., 2003). However, the interactions with several such E3 ligases (DWA1, DWA2 and KEG1) are rather weak and loss of function mutations for these ligases do not block the ABA resistance conferred by AFP2 overexpression (Lynch et al., 2022). Furthermore, tests of the timecourse of germination show that germination mostly precedes degradation of ABI5 (Lynch et al., 2022) and COP1 promotes ABI5 accumulation in the dark (Peng et al., 2022).

Outside of the conserved domains, much of the AFP sequences are predicted to be intrinsically disordered (Lynch et al., 2017; Varadi et al., 2023) (Supp. Fig.1), raising the possibility that the AFPs can assume a variety of structures due to post-translational modifications, consistent with the diversity of their observed protein interactions. Furthermore, much recent work highlights the tendency of intrinsically disordered regions to drive phase transitions forming bimolecular condensates (Liu et al., 2023). The current work reports interactions between AFPs and the PP2Cs, SnRK2s and MPKs previously shown to regulate ABA response, and addresses the functional significance of these interactions. Additional interactions with ABA receptors and 14-3-3 proteins may affect the activity of signaling complexes containing some of these factors.

## RESULTS

### Interactions between AFPs and components of the ABA core signaling pathway

Our early studies of the AFPs revealed no obvious biochemical function, so we used AFP2 as bait in a yeast two-hybrid screen to identify additional interacting partners that might provide clues regarding function. This screen produced a diverse set of potential interactors including multiple ABI5/ABF/AREB clade bZIP proteins, ABI clade protein phosphatases (PP2Cs), and a variety of kinases. Further specific pairwise yeast two-hybrid assays, expanding this study to include all four members of the AFP family and seven members of the ABI PP2C clade, showed strong interactions between all AFPs and AHG1 and AHG3. When the AFPs were present as Binding Domain (BD)-fusions, additional interactions were limited to either AFP1 or AFP2 for only one other PP2C tested, ABI1 (Fig.1). Interactions mapped to the C domain of AFP1, but to several regions of AFP2.

**Fig. 1.**
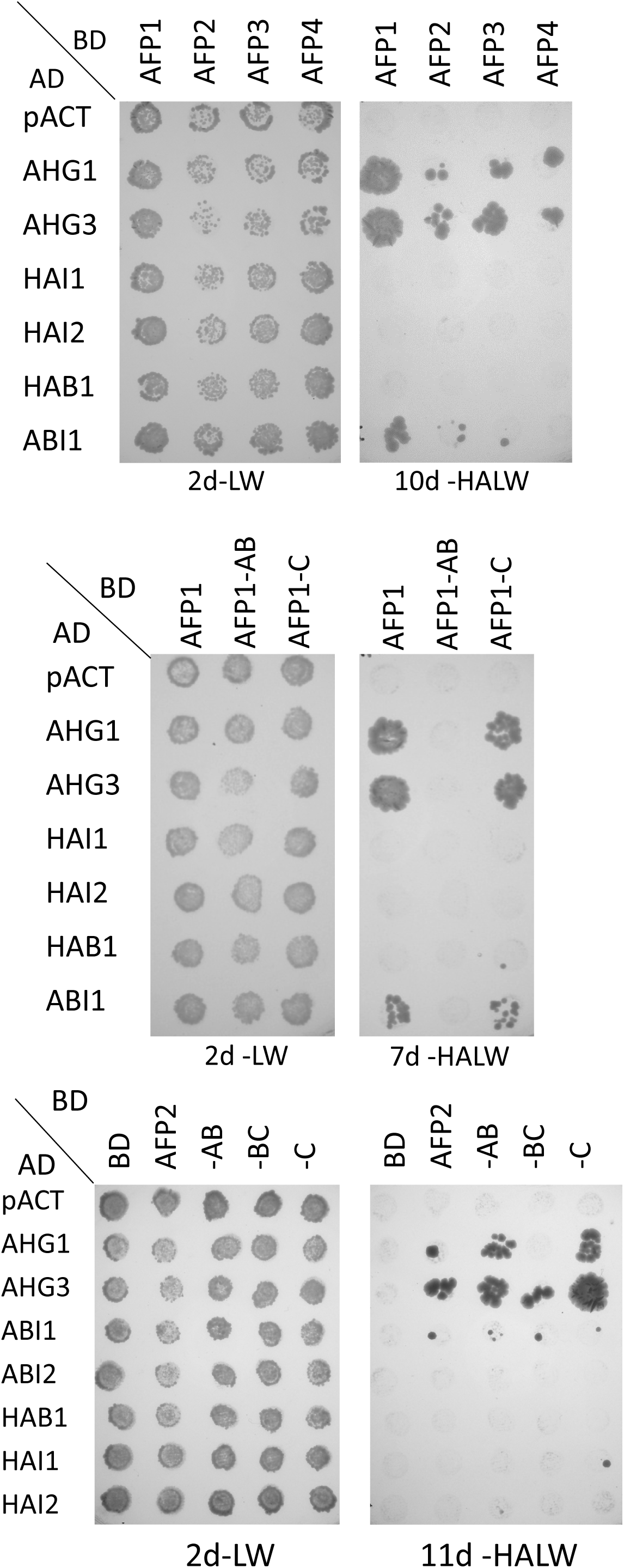
Interactions between AFPs and PP2Cs detected by yeast two hybrid assays. Fusions were combined by matings between Y187 carrying AD fusions and PJ69-4a carrying BD fusions. After overnight incubation on YPD media, yeast were replica-plated onto selective media lacking leu and trp (-LW) to maintain the AD-and BD-fusion plasmids, and lacking histidine and adenine (-HALW) to score interactions between the fusion proteins.

Tests of interactions with kinases showed all AFPs except AFP4 interacted with SnRK2.3, SnRK2.6, MPK3, MPK4, and MPK7, but not with SnRK2.10 (Fig.2). The BD-MPK6 fusion conferred growth without requiring an AD-interaction partner, but reporter activation was again enhanced by interaction with AD-fusions to AFP1,2 and 3 (Supp. Fig. 2). Surprisingly, activity was impaired when BD-MPK6 was co-expressed with AFP4 or AFP2-C. Only AFP2 interacted with SnRK2.2 in this assay. Interactions with subdomains were very weak, suggesting that regions across the entire protein were required for these interactions. Although several D clade MPKs are highly co-expressed with the AFPs in maturing, dry and germinating seeds (Supp.Fig 3), only MPK17 and MPK20 interacted with the AFPs in yeast, and the interactions were relatively weak.

**Fig. 2.**
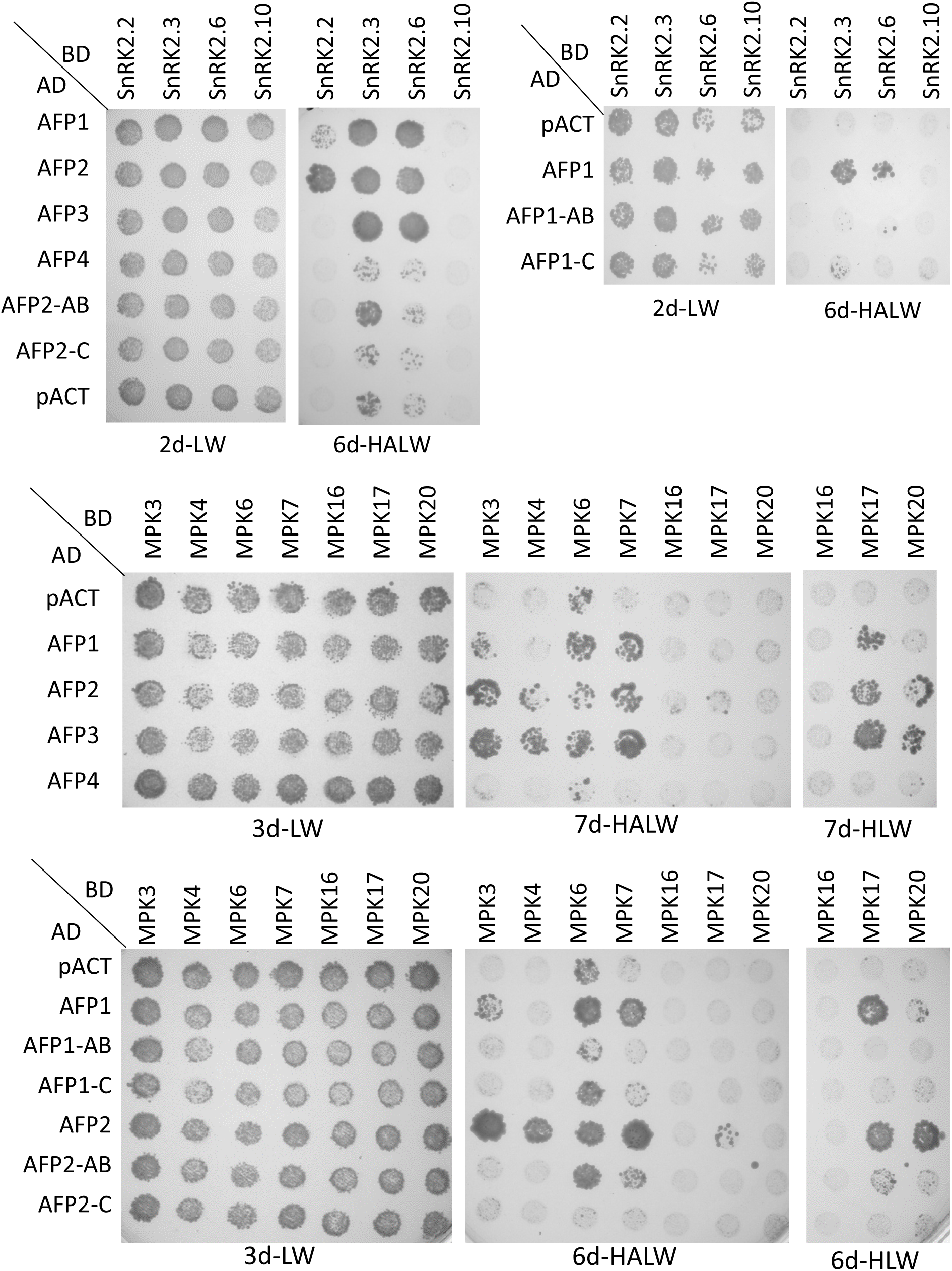
Interactions between AFPs and kinases detected by yeast two hybrid assays. The indicated fusions were combined and interactions were scored as described in Fig. 1.

In contrast to the strong interactions between AFPs and bZIPs, PP2Cs and kinases, relatively weak interactions were observed between AFPs and a few members of the PYR/PYL/RCAR family of ABA receptors: PYLs 2, 7, 9, 11 and 12 (Fig. 3A, Supp Fig. 4). All of these are expressed in maturing, dry and/or imbibing seeds, coinciding with peak expression of AFP1, AFP2, and ABI5 (Supp. Fig. 5).

**Fig. 3.**
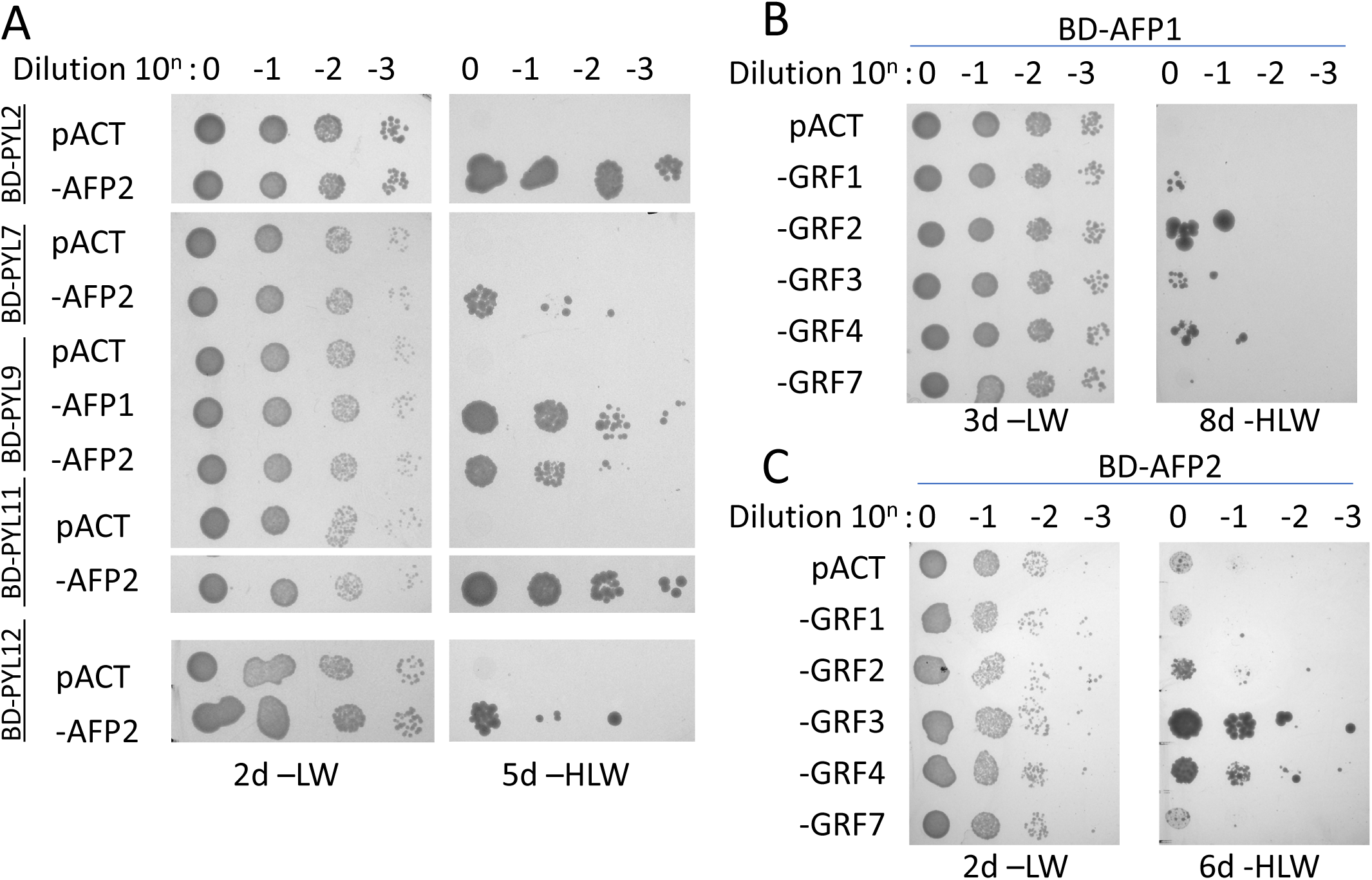
Interactions between AFPs, receptors and GRFs detected by yeast two hybrid assays. Following matings, the indicated diploid lines were grown overnight in media lacking leu and trp. After measuring the OD600, cultures were diluted to the same concentration, then serially diluted 10-fold three times. The diluted cultures were then replica-spotted onto selective media. The -LW plate serves as a control for accuracy of the dilutions. Growth on the -HLW plate requires interaction between the AD- and BD-fusions to complement the *his3-200* allele of these yeast.

Furthermore, PYLs 11 and 12 promote ABI5-mediated ABA response during germination and their expression is regulated by ABI5 (Zhao et al., 2020), and PYLs 7 and 9 are members of the small subset of receptors that can interact with AHG1 (Tischer et al., 2017). This result suggests that several AFPs can interact with all components of the ABA core signaling pathway.

Many phosphorylation-dependent protein activities, including hormone signaling, are regulated by 14-3-3 proteins that bind to phosphorylated Ser or Thr residues within specific contexts (Camoni et al., 2018). The Arabidopsis genome encodes thirteen expressed 14-3-3 proteins, designated GENERAL REGULATORY FACTORS(GRFs), many of which are also expressed in maturing, dry and germinating seeds (Supp Fig 5). Previous studies in multiple cereal grains and Arabidopsis have shown a role for 14-3-3 proteins in ABA-regulated transcriptional complexes, including direct interactions with members of the ABI5/ABF clade of bZIPs (Jaspert et al., 2011). We initially tested the Arabidopsis 14-3-3 proteins with the highest homology to the barley and rice proteins shown to interact with HvABI5 and HvABF1 (Schoonheim et al., 2007), and that are expressed in dry, imbibed or germinating seeds: GRF1 (14-3-3 chi), GRF2 (14-3-3 omega), GRF3 (14-3-3 psi), GRF4 (14-3-3 phi), and GRF7 (14-3-3 nu). AD fusions with GRF1,2,3, and 4 all interacted weakly with a BD-AFP1 fusion, but only the AD-GRF3 and -GRF4 fusions interacted above background with BD-AFP2 (Fig. 3BC). Interaction with ABI5 was substantially stronger (Supp Fig 6). Consistent with the importance of phosphorylation for 14-3-3 binding, a BD-fusion to AFP2 with alanine substitution mutations in 2 binding sites predicted by 14-3-3-Pred (https://www.compbio.dundee.ac.uk/1433pred/)(Madeira et al., 2015) had greatly reduced interactions with the GRFs (Supp Fig 6). In addition to their relatively strong interactions with AFP1 and AFP2, transcripts for both GRF3 and GRF4 are present at similar levels to those of these AFPs in dry and imbibed seeds (Supp. Fig. 3), so our further studies focused on interactions with these family members.

We previously found that AFP1 and AFP2 could form homo- and heterodimers in yeast two-hybrid and BiFC assays (Lynch et al., 2017). Attempts to map the interacting domains by further yeast two-hybrid assays showed that the AB domain was sufficient for interactions with full-length AFP2, but not with itself, implying that multiple domains are required for interaction (Supp. Fig. 7A). Consistent with this, Alphafold modeling suggests that the most likely interface between AFP2 subunits includes residues present in the B and C domains (Supp. Fig. 7B).

The interactions between AFP1, AFP2, PP2Cs, kinases, receptors and GRFs were further tested by BiFC assays, using split YFP fusions in transiently transformed *N. benthamiana*. These studies showed interactions between both AFPs and SnRK2.3, SnRK2.6, MPK3, MPK4, MPK6, AHG1, AHG3, GRF3, GRF4, PYL2 and PYL9, confirming the yeast two-hybrid results, but interactions with GRF7 and AHG3 were much weaker in these assays (Fig.4). AFP2 also interacted with MPK7 in this assay. Although none of the 14-3-3 proteins showed any specific interaction with PP2Cs or a SnRK that interacts with AFP2 in the yeast two-hybrid assay, GRF4 interacted strongly with MPK3 and SnRK2.6, and very weakly with AHG1, in BiFC assays (Supp Fig.6). Further mapping of the BiFC interactions between domains of the AFPs with AHG1 and SnRK2.3 showed that all AFP domains interacted with AHG1 (Supp Fig 8) (Erickson McNally, 2016). In contrast, SnRK2.3 interacted with only the B and C domains of AFP2, and the A and B domains of AFP1.

The BiFC assays also showed some differences in localization. Fluorescence was generally diffuse throughout the nuclei when AFPs and kinases were co-expressed, but localized to nuclear punctae when AFPs and AHG1 were combined (FIg 4 and Supp Fig. 8). Although GRF4 was distributed throughout the cell when forming homodimers, or heterodimers with GRF3 and GRF7, the GRF interactions with AFPs were mostly limited to nuclei (Fig.4).

**Fig. 4.**
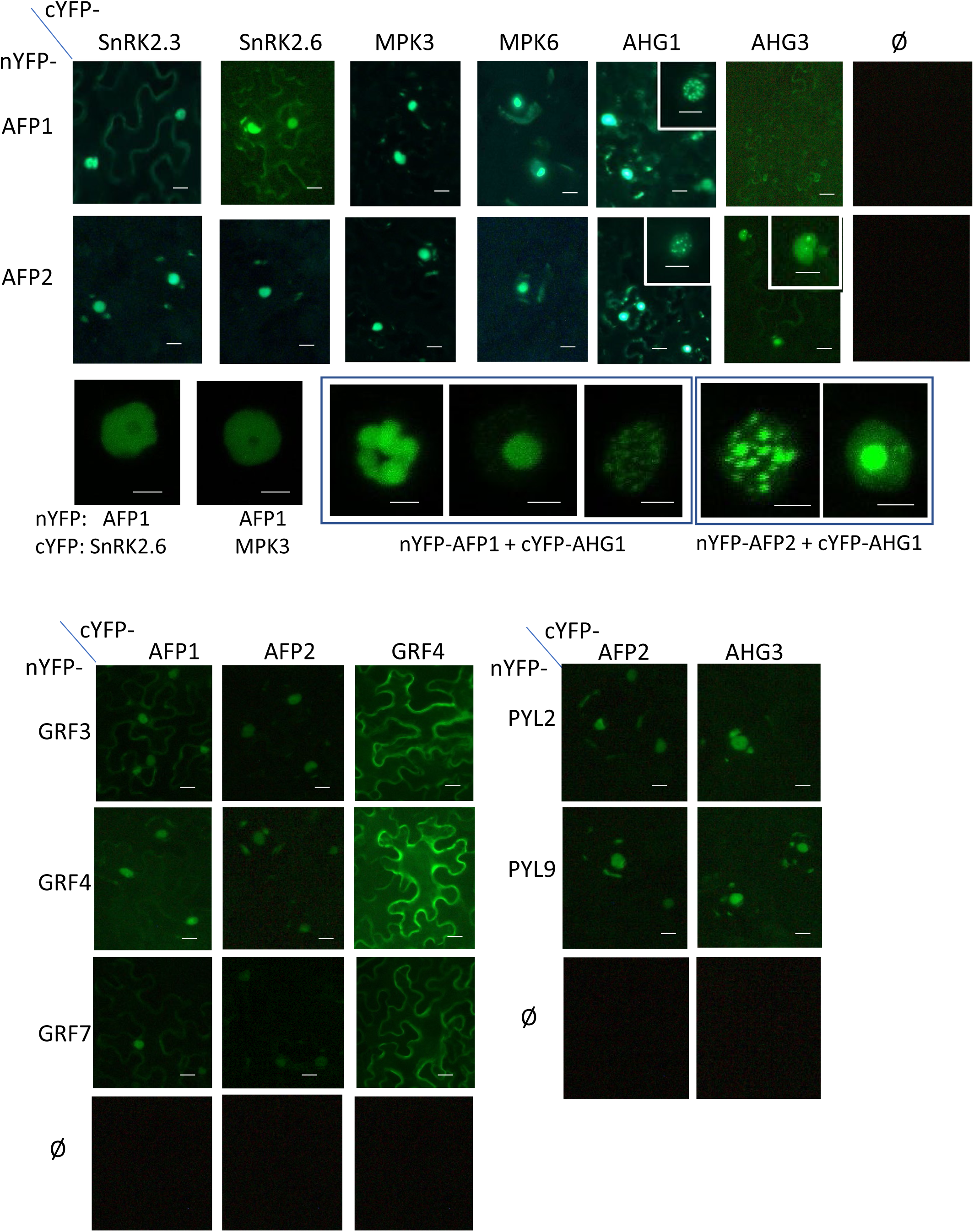
Interactions between AFPs and SnRK2s, PP2Cs, GRFs and receptors detected by split YFP assays. Low magnification images were obtained with an Infinity camera on an Olympus AX70 epifluorescence microscope at 100x magnification. Insets are 400x magnification. High magnification images of individual nuclei were obtained with a Leica SP8 Resonant Scanning Confocal microscope. Scale bars = 10 μm for interactions with kinases, 5 μm for interactions with AHG1

### AFPs are substrates for SnRK2s and PP2Cs

We had previously observed that YFP-AFP2 fusions migrate as doublets and occasionally even triplets on SDS-PAGE, suggesting the possibility of post-translational modifications (Erickson McNally, 2016). These doublets are especially pronounced in any fusions containing the B domain of AFP2 (Fig. 5A). To determine whether the observed interactions between AFP2 and these kinases and phosphatases reflected a substrate role of AFP2, these proteins were transiently co-expressed in *N. benthamiana*. Co-expression with SnRK2.3 shifted the YFP-AFP2-AB fusion to predominantly the slower mobility form, whereas co-expression with AHG1 or AHG3 resulted in a shift toward the higher mobility form (Fig.5B). YFP-AFP2-AB affinity purified with GFP-trap beads cross-reacted with an anti-phosphoserine antibody, and this reaction was enhanced in the slower migrating form, confirming that the altered mobility was due at least in part to phosphorylation (Fig.5C).

**Fig. 5.**
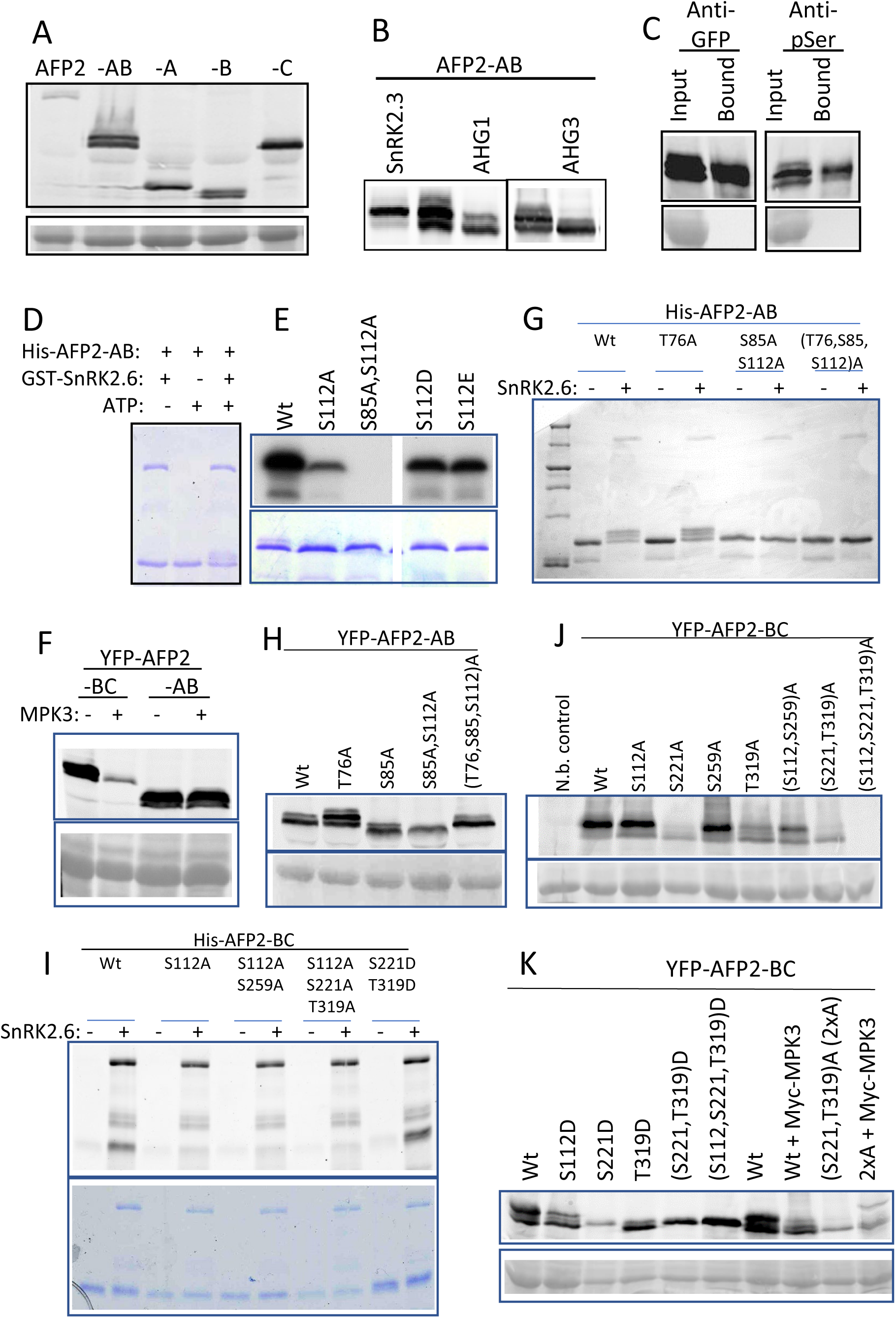
AFP2 is a substrate for both specific kinases and PP2Cs. (A) Immunoblot of protein extracts from *N. benthamiana* leaves with transiently expressed full-length or subdomains of AFP2 as YFP fusions, detected by anti-GFP antibody; lower panel is Ponceau stain focused on Rubisco as loading control. (B) Immunoblot of YFP-AFP2-AB fusion transiently co-expressed with fusions to either SnRK2.3 or AHG1. (C) Transiently expressed YFP-AFP2-AB was immunoprecipitated using GFP-Trap agarose beads; upper panels are immunoblots with indicated antibodies, lower panels are Ponceau stain. *In vitro* kinase reactions with GST-SnRK2.6 and wt or phosphomutant His-fusions of AFP2-AB resulting in reduced mobility of the fusion protein (D) detected by Coomassie stain, (E) incorporation of radioactivity, lower panels are Coomassie stain of gel used in autoradiograph, or (G) reactivity with ProQDiamond stain. Immunoblots of protein extracts from leaves transiently expressing YFP fusions of (F) the wild-type AB and BC domains of AFP2 with Myc-MPK3, (H) wild-type and phosphomutant AFP2-AB domain fusions, (J) wild type and phosphomutant AFP2-BC domain fusions, and (K) wild type and phosphomimetic AFP2-BC domain fusions, including some co-expressed with Myc-MPK3. YFP fusions were detected by anti-GFP antibody, lower panels are Ponceau stain. (I) *In vitro* kinase reactions with GST-SnRK2.6 and wt or mutant His6 fusions of AFP2-BC, viewed with ProQDiamond stain, lower panel is subsequent Coomassie stain of same gel.

AFP2 contains 40 serines and 22 threonines, constituting nearly 18% of the total amino acids in AFP2, many of which are conserved between AFP1 and AFP2 (Supp. Fig. 9). Of these, 17 serines and 11 threonines were predicted to be phosphorylated by the PhosPhAt database (Zulawski et al., 2012), but only 9 serines and 1 threonine are predicted at high stringency by http://musite.sourceforge.net/ (Table 1). PhosPhAt lists only three documented phosphorylated residues, following ionizing radiation stress and dependent on the ataxia telangiectasia-mutated (ATM) and Ataxia telangiectasia-mutated and Rad3-related (ATR) kinases: T76, S232 and S234 (Roitinger et al., 2015). Of these, only T76 was predicted by both programs. Based on the consensus target sites of R-X-X-S and S-D for ABA-induced phosphorylation by SnRK2s (Wang et al., 2013), potential targets in AFP2 are S85, S112, S150, S187, and S259. Of these, only S85 and S112 are conserved between AFP1 and AFP2. Similarly, two consensus sequences, P-X-S/T-P and S/T-P, are highly abundant in protein substrates of MAPKs (Seger and Krebs, 1995) such that potential targets of MAPKs are T76, S221 and T319 of AFP2, all of which are conserved between AFP1 and AFP2. Three consensus motifs have been proposed for 14-3-3 target binding sites, two of which are present in AFP2: S112 fits the mode I consensus and both S346 and T347 fit the C-terminal mode III consensus (reviewed in Camoni et al., 2018). The 14-3-3 Pred program scores S112 and S85 as highly likely binding sites, with relatively moderate scores for T16, S38, and S259.

**Table 1.** AFP2 predicted phosphorylation sites.

| Residue | NetPhos<br>2.0<br>score | PhosPhAt<br>score | MUsite<br>score | SnRK2<br>target | MPK<br>target | in vitro<br>phosphorylated |
| --- | --- | --- | --- | --- | --- | --- |
| Y28 | 0.315 | 1.59 |  |  |  |  |
| S38 | 0.02 | 0.11 | 0.487 |  |  |  |
| T51 | 0.781 | 0.21 | 0.212 |  |  |  |
| T76 | 0.973 | 1.1 | 0.841 |  | T-P |  |
| S83 | 0.971 |  | 0.769 |  |  |  |
| S84 | 0.947 |  | 0.787 |  |  |  |
| S85 | 0.997 | 0.81 | 0.881 | R-S-S-S |  | Yes |
| T89 | 0.014 | 0.84 | 0.292 |  |  |  |
| Y103 | 0.622 | 0.97 |  |  |  |  |
| T104 | 0.041 | 0.19 | 0.196 |  |  |  |
| T111 | 0.9 |  | 0.21 |  |  |  |
| S112 | 0.321 | 0.37 | 0.77 | R-T-T-S |  | Yes! |
| S129 | 0.988 |  | 0.469 |  |  |  |
| S150 | 0.753 | 0.61 | 0.46 | S-D |  |  |
| T154 | 0.084 | 0.14 | 0.383 |  |  |  |
| S156 | 0.966 | 0.45 | 0.737 |  |  |  |
| T164 | 0.471 | 0.75 | 0.113 |  |  |  |
| S187 | 0.179 | 0.67 | 0.578 | S-D |  |  |
| T193 | 0.137 | 0.91 | 0.298 |  |  |  |
| S197 | 0.323 | 0.44 | 0.277 | G-S? |  |  |
| S198 | 0.725 | 0.52 | 0.227 |  |  |  |
| S199 | 0.063 | 0.21 | 0.154 |  |  |  |
| S200 | 0.992 | 0.65 | 0.539 |  |  |  |
| S202 | 0.996 |  | 0.274 |  |  |  |
| S213 | 0.572 | 0.03 | 0.249 |  |  |  |
| S215 | 0.916 | 0.8 | 0.402 |  |  |  |
| S221 | 0.772 | 1.13 | 0.654 |  | S-P |  |
| S232 | 0.844 | 0.46 | 0.47 |  |  |  |
| S234 | 0.745 | 0.56 | 0.536 | G-S? |  |  |
| T243 | 0.046 | 0.21 | 0.137 |  |  |  |
| S259 | 0.985 | 1.28 | 0.869 | R-V-R-S |  |  |
| S262 | 0.763 | 0.63 | 0.803 |  |  |  |
| S273 | 0.044 | 0.18 | 0.348 |  |  |  |
| S284 | 0.303 | 0.34 | 0.103 |  |  |  |
| Y299 | 0.137 | 0.46 |  |  |  |  |
| T319 | 0.94 | 0.27 | 0.247 |  | T-P |  |
| T343 | 0.689 |  | 0.173 |  |  |  |

We took two approaches to identifying actual phosphorylation sites: *in vitro* phosphorylation with purified recombinant proteins, and site-directed mutagenesis of predicted phosphorylation sites. We focused on specific target sites of SnRK2s by incubating recombinant His6-AFP2-AB (aa1-149) fusions with GST-SnRK2.6 *in vitro*, resulting in a shift of the His-AFP2-AB fusion to a higher apparent molecular weight (Fig. 5D). Mass spectrometric analysis of this slower mobility form identified S112 as a phosphorylated residue (Supp Fig 10). *In vitro* kinase assays including gamma-labeled ATP, comparing wild-type to either S112A or phosphomimetic S112E or S112D mutants of His6-AFP2-AB, showed that phosphorylation by SnRK2.6 was still possible until the S85 was also converted to alanine (Fig. 5E). The phosphomimetic mutants appeared to have an intermediate degree of phosphorylation between the wild type and S112A forms, suggesting that true phosphorylation at S112 affected the likelihood of additional phosphorylation. In contrast, a His6-AFP2-C fusion (aa-149-348) was not phosphorylated by SnRK2.6 (Supp Fig 11A).

Immunoblot analyses of possible *in vivo* phosphorylation by Myc-MPK3 showed either no shift in mobility for the YFP-AFP2-AB domain fusion or an apparent compression to the higher mobility form for the YFP-AFP2-BC domain fusion (Fig.5F). Surprisingly, although neither the S85A, S112A double mutant nor the T76A, S85A, S112A triple mutant of the His6-AFP2-AB domain fusion could be phosphorylated by SnRK2.6 *in vitro* (Fig 5G), only the double mutant ran as a single high mobility form when transiently expressed as a YFP-fusion *in planta* (Fig.5H). The triple mutant ran at the same mobility as wild-type when expressed as a YFP fusion *in planta*. and the T76A mutant of the His6-AFP2-AB domain fusion showed a greater shift toward lower mobility forms than the wild-type fusion (Fig. 5H). However, similar to the wild-type fusion, its mobility was not altered by co-expression with Myc-MPK3 (Supp. Fig. 11B).

*In vitro* kinase assays with His6-AFP2-BC domain fusions showed that an S112A substitution greatly reduced phosphorylation of this domain by SnRK2.6 (Fig.5I). Similarly, S112A mutations blocked *in planta* phosphorylation of YFP-AFP2-BC fusions by SnRK2.3 when transiently co-expressed in *N. benthamiana* (Supp Fig. 12A). Although phosphorylation of S112A in the *in vitro* reactions was not affected by phosphomimetic mutations in S221 and T319, these mutations reduced mobility of the fusion (Fig. 5I).

Transiently expressed YFP-AFP2-BC mutant fusions converting S112, S259, S221 or T319 to alanine all still appeared as doublets in immunoblot analysis, as did double mutants combining predicted SnRK2 targets (S112, S259)A or potential MPK targets (S221, T319)A (Fig. 5J). However, both single and double mutants affecting S221 and T319 reduced the shift toward lower mobility, and both double mutants were under-accumulated relative to the wild type. The triple mutant (S112,S221,T319)A fusion was rarely detectable. Unlike the alanine substitution mutants, YFP-AFP2-BC fusions with phosphomimetic mutations converting S221 alone or in combination with S112 and T319 to aspartic acid all resulted in compression to a single intermediate mobility form (Fig.5K). Co-expression with MPK3 also converts most of the wild type AFP2-BC fusion toward higher mobility forms, but has little effect on the (S221,T319)A mutant which already displays a higher mobility than the wild type (Fig.5F,K). In contrast to the higher mobility on Laemmli SDS-PAGE of AFP2-BC co-infiltrated with MPK3, analysis on Phos-tag gels showed a shift toward decreased mobility, reflecting increased phosphorylation (Supp Fig 12B). The S221A mutation alone blocked the phosphorylation seen in the wild-type BC domain, but a phosphorylated form reappeared in the (S221,T319)A mutant. Although the (S221,T319)A mutant migrated similarly to the wild-type BC domain, co-expression with MPK3 resulted in the appearance of a novel phosphorylated form, visible in both gel systems, suggesting that loss of both “consensus” MPK targets led to phosphorylation of an as-yet unknown alternative residue, possibly by an unidentified endogenous kinase.

In contrast to the AB and BC domain fusions, full-length YFP-AFP2 transiently co-expressed with Myc-AHG1 in *N. benthamiana* shows as much or more phosphorylation than YFP-AFP2 alone, even when the SnRK2 and potential MPK target residues have been converted to alanines, again suggesting that alternative residues are being modified (Supp Fig.12D). Surprisingly, YFP-AFP2 co-expressed with Myc-SnRK2.3 appeared less phosphorylated than YFP-AFP2 alone. However, alanine substitutions in the SnRK target residues (S85 and S112) resulted in the appearance of novel bands, reflecting both less and more phosphorylation, but including the additional S259A mutation fully blocked SnRK2-induced phosphorylation (Supp Fig.12D). Alanine substitutions in potential MPK target residues (S221 and T319) also blocked phosphorylation by SnRK2.3. Collectively, these results suggest that modification of the primary SnRK2 or MPK targets affects accessibility of the SnRK2 target sites and may lead to phosphorylation of novel sites.

### ABA effects on AFP2 phosphorylation state

Given that ABA activates the group 3 SnRK2s, we tested whether these mutations affected any post-translational modifications induced by ABA treatment. Transiently expressed wild type YFP-AFP2-AB domain fusions shifted from roughly 40% slower mobility forms in control conditions to 60% slower mobility forms after 4 hrs exposure to 30 μM ABA (Fig. 6A). In contrast, the S112A mutation alone, or in combination with T76A and S85A (“3xA”), blocked this shift (Fig. 6A). Similarly, the highest mobility form of the YFP-AFP2-BC domain fusion shifted to an intermediate slower mobility following ABA treatment and this was blocked in the (S112,S259)A mutant. In contrast, the (S221,T319)A mutant of the BC domain continued to show a shift of the highest mobility form in response to ABA, but had greatly reduced expression of the slowest mobility form under any condition.

**Fig. 6.**
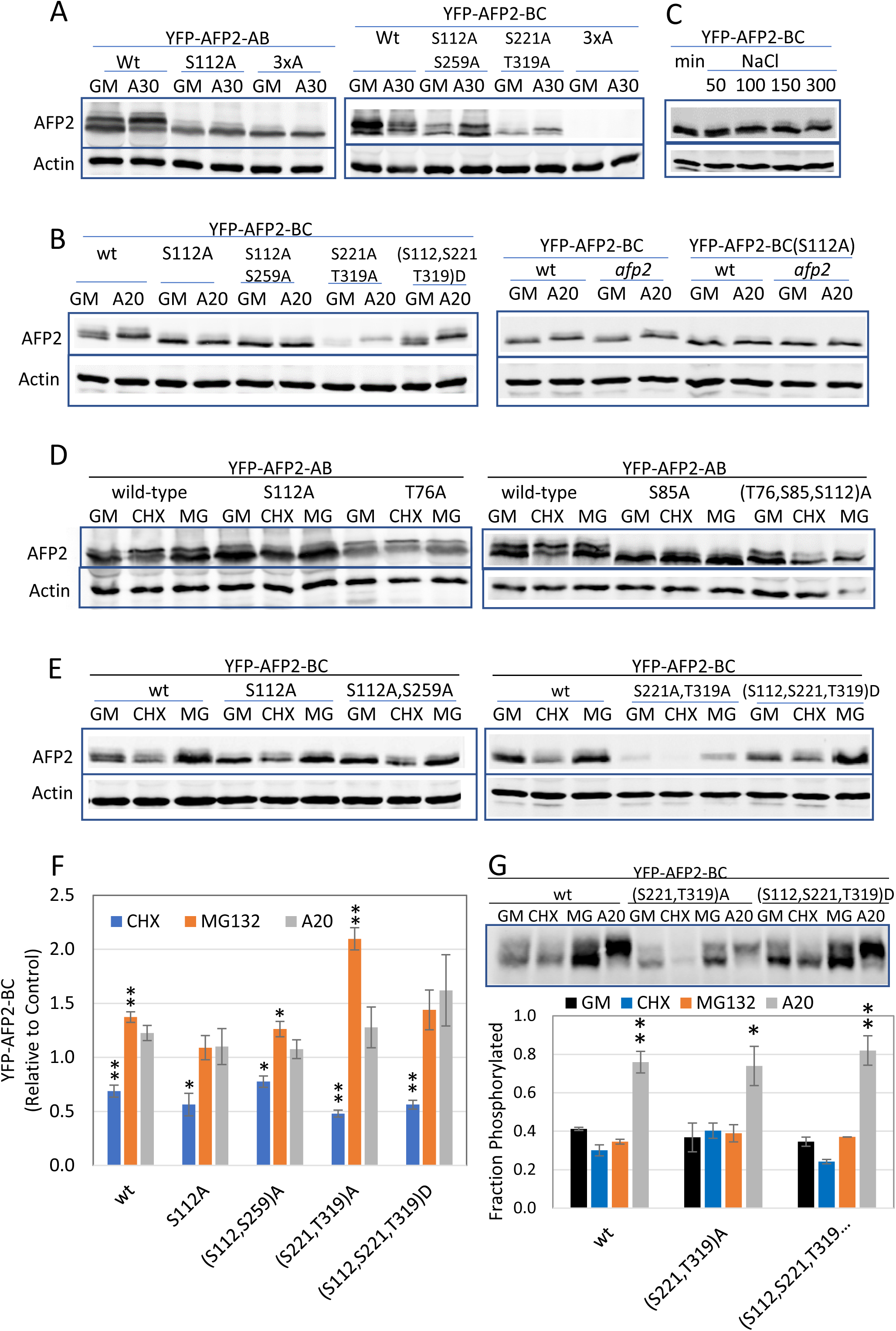
*In vivo* expression assays with mutant AFPs testing relative significance of specific residues for ABA/stress-induced modification and stability. Transient expression in *N.benthamiana*, followed by 4 hrs incubation in GM +/- 30 μM ABA (A). Stable *A. thaliana* transgenic seedlings grown 5d on GM, then incubated in GM +/- 20 μM ABA for 4.5 hrs (B) or min +/- NaCl (50 - 300 mM) for 2.75 hr (C). Transient expression of YFP-AFP2-AB fusions in *N.benthamiana*, followed by 6 hrs incubation in GM +/- 100 μM CHX or MG132 (D). Stable *A. thaliana* transgenic seedlings grown 5d on GM, then incubated in GM +/- CHX or MG132 for 6 Hrs (E).

In stable transgenic Arabidopsis lines carrying the wild-type YFP-AFP2-BC domain fusion, this shift in mobility was induced by exposure to as little as 1 μM ABA for 6 hrs or by 1 hr of treatment with 20 μM ABA (Supp Fig 13AB). Similar to the transient expression assays, only those lines with the S112A mutation blocked the shift to slower mobility (Fig. 6B), suggesting that this residue is modified in response to ABA treatment. Phos-tag gel analysis showed that the ABA-induced shift in mobility correlated with S112-dependent changes in phosphorylation (Supp Fig 12C), but surprisingly, “phosphomimetic” Asp substitutions at S112, S221, and T319 (“3xD”) did not prevent phosphorylation of this fusion. A slight mobility shift of the YFP-AFP2-BC domain fusion was also seen in response to salinity stress imposed by at least 100 mM NaCl (Fig.6C).

Although transiently expressed full-length YFP-AFP2 fusions showed no difference in phosphorylation between control and ABA treatments, there was generally a higher degree of phosphorylation in proteins with more ala substitutions, again consistent with modifications at novel sites (Fig.7A). In contrast, transgenic lines carrying the full-length YFP-AFP2 fusion showed an ABA-induced shift toward slower mobility/greater phosphorylation of the fusion (Fig.7B). The stable transgenic lines with (S85,S112)A substitutions showed no change in fusion mobility in response to ABA, whereas the phosphomimic mutations (S85,S112)E displayed a shift toward slower mobility on control media.

**Fig. 7.**
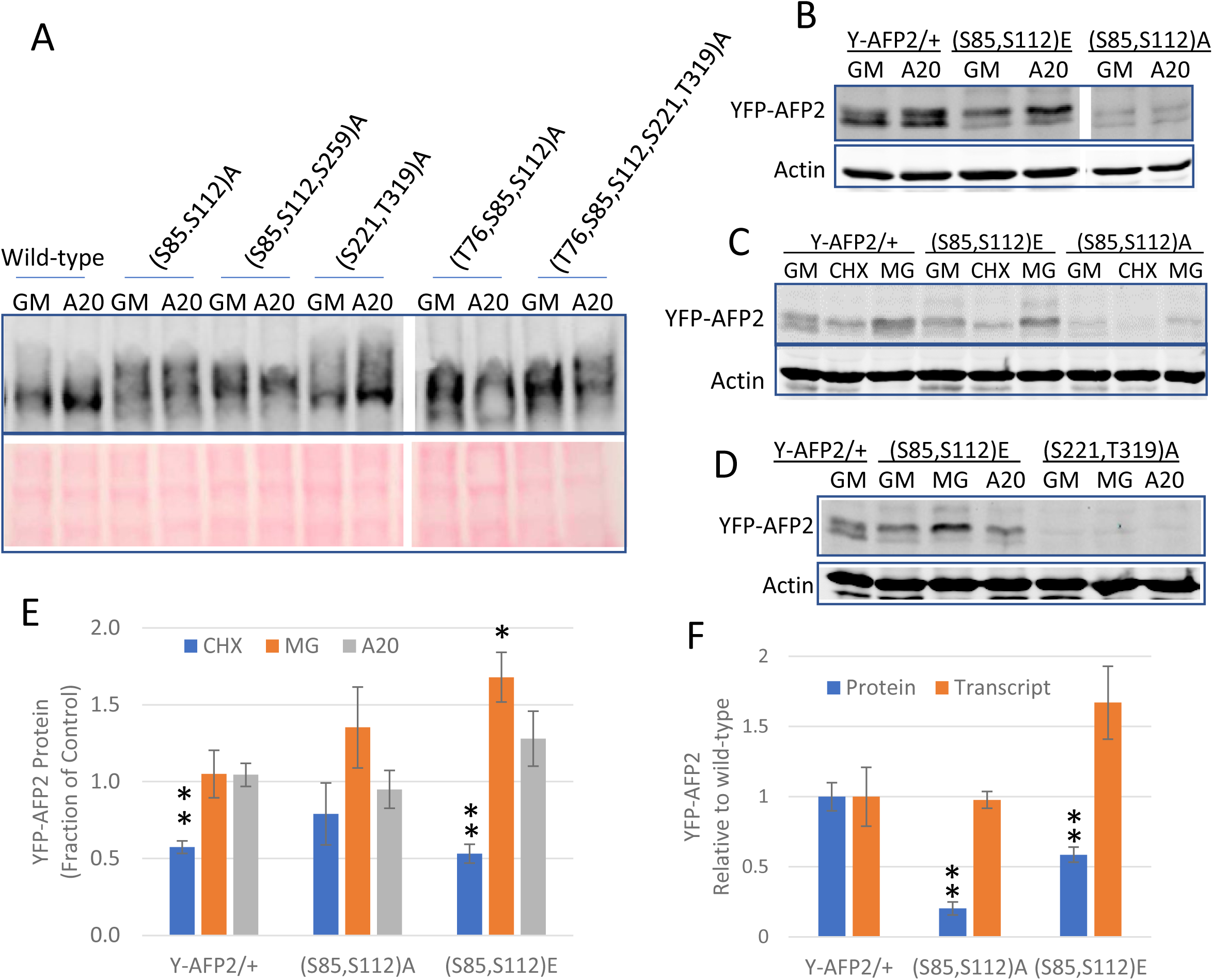
Expression assays with full-length AFP2 fusions testing relative significance of specific residues for ABA-induced modification and stability. (A) Immunoblots of protein extracts from *N. benthamiana* leaves transiently expressing YFP fusions of full-length AFP2 with the indicated substitutions, following 6 h exposure to control media (GM) +/-20 μM ABA (A20), resolved on a Phos-Tag gel. YFP-AFP2 fusions were detected with anti-GFP antibody; lower panel is Ponceau stain. (B-D) Immunoblots of protein extracts from stable transgenic Arabidopsis 5d old seedlings with the indicated YFP-AFP2 fusions, following 6h exposure to GM, supplemented with 20 μM ABA (A20) or 100 μM cycloheximide (CHX) or MG132 (MG). Proteins were resolved on 10% SDS-PAGE. (E) Accumulation of fusion protein following treatment with CHX, MG132 or ABA, normalized relative to actin, expressed relative to level of that fusion in seedlings incubated on GM, as shown in Figs. 7B-D. (F) Relative protein and transcript levels of wild-type and mutant YFP-AFP2 fusions in 5d old seedlings of stable transgenic lines. YFP-AFP2 protein levels were normalized relative to actin. Transcript levels were normalized relative to the geometric mean of PP2AA3 (At1g13320) and AP2M (At5g46630) of Arabidopsis. Both are expressed relative to the average value for the wild-type construct. Data shown in E & F is average of at least triplicate samples, error bars represent S.E.** and * indicate statistically different from control (P <0.01 and P<0.05, respectively, based on two-tailed Student’s t-test)

Comparison of transcript levels for wild-type vs. mutant fusions in tissue harvested in parallel to that used for protein extracts showed that levels were somewhat variable for all constructs, as expected for a transient system, but the reductions in protein levels of the mutant forms were far greater than any changes in transcript levels (Supp Fig 14). Although some transiently expressed transgene transcripts were two-to three-fold lower for the mutant constructs, this was not sufficient to explain the at least ten-fold decreases in YFP-AFP2-AB protein or the five-to ten-fold decreases in YFP-AFP2-BC protein fusions (Supp. Fig. 14C). Stable transformants carrying the (S221,T319)A mutant YFP-AFP2-BC fusion also showed at least five-fold less protein despite a similar transcript level as the wild-type fusion. Similarly, the YFP-AFP2-BC(S112A) mutant form showed a roughly four-fold reduction in protein:transcript in stable transformants (Supp. Fig. 14E). All of these results are consistent with possible defects in stability or translation of the fusions. Protein levels were greatly reduced for full-length YFP-AFP2 fusions with ala substitutions in stable transgenic lines, but transcript levels were similar for wild type and the (S85,S112)A variant (Fig. 7).

### Effects of phosphorylation on AFP2 stability

To further address the functional relevance of these differences in phosphorylation, we tested stability and potential proteasomal degradation by assaying accumulation following treatment with either the translation inhibitor cycloheximide (CHX) or the proteasome inhibitor MG132. In transient expression assays, the higher mobility form of the AB domain decreased from roughly 58% in control conditions to 50% of the total in CHX, but the high mobility form was stabilized by MG132 (Fig.6D), consistent with proteasomal degradation of the high mobility/unphosphorylated form. Accumulation of the low mobility form was still seen in the T76A mutant, but was reduced in all the mutant fusions affecting SnRK2 targets: S85A, S112A, (S85,S112)A and the (T76,S85,S112)A triple mutant, consistent with a role for SnRK2-mediated phosphorylation in regulating protein stability. The S85A mutant appeared to have an even higher mobility than the wild-type fusion forms (Fig.5H,6D).

The YFP-AFP2-BC domain fusions were tested in both transient and stably transformed tissue. All stable transformants showed reduced fusion accumulation following CHX treatment and most increased accumulation in MG132-treated seedlings (Fig. 6E,F). The fusion combining S221A and T319A mutations had severely decreased accumulation in both stable and transiently transformed tissue (Fig. 6E,F and Supp.Fig. 14DE). ABA treatment primarily stabilized accumulation of the phosphorylated form (Fig 6G). Although the double mutant proteins were stabilized by MG132 treatment, this treatment was not sufficient to increase accumulation of the (S112, S221, T319)A (3xA) mutant in transiently transformed tissue (Supp.Fig. 15). Furthermore, the limited YFP-AFP2-BC(3xA) fusion visible was present in only a small fraction of cells, mostly in small punctae throughout the cytoplasm, not concentrated in the nuclei as for all other YFP-AFP2 fusion constructs (Supp. Fig. 15C).

In stable transformants with full-length wild-type YFP-AFP2, the phosphorylated form was more stable than the faster-migrating form (Fig.7C). Consistent with this, lines with either alanine or phosphomimetic substitutions affecting S112 have substantially lower fusion protein levels, despite transcript levels that are similar or higher than those for the wild-type fusion (Fig. 7CF). All of these full-length YFP-AFP2 fusions are proteasomally degraded, but the effects of CHX and MG132 are more limited for those with ala substitutions (Fig. 7CE, Supp. Fig. 17F). In contrast, the only stable lines expressing the (S221,T319)A variant had extremely low fusion protein accumulation, which was not increased by MG132 treatment (Fig.7D) and reflected very low transcript levels (less than 0.1% of that for the wild-type fusion transcript).

### Effects of phosphorylation on AFP2 localization

Having observed that co-expression of either nYFP-AFP fusions with cYFP-AHG1 or YFP-AFP2 with Myc-AHG1 led to formation of nuclear punctae we tested whether mutations preventing phosphorylation would lead to similar changes in localization in transient expression assays. Whereas full-length YFP-AFP2 fusions appear mostly diffuse through nuclei, but generally excluded from nucleoli, YFP-AFP2-AB domain fusions display as dense bundles of foci (Fig. 8). YFP-AFP2-AB(S112A) and (3xA) mutants appear to form fewer foci. Although not uniform across the leaf, the two double mutants of the YFP-AFP2-BC domain fusions tend toward more discrete areas of accumulation, appearing either more excluded from some regions (S112A,S259A) or more focused in a few foci (S221A,T319A), consistent with effects of phosphorylation state on changes in aggregation state.

**Fig. 8.**
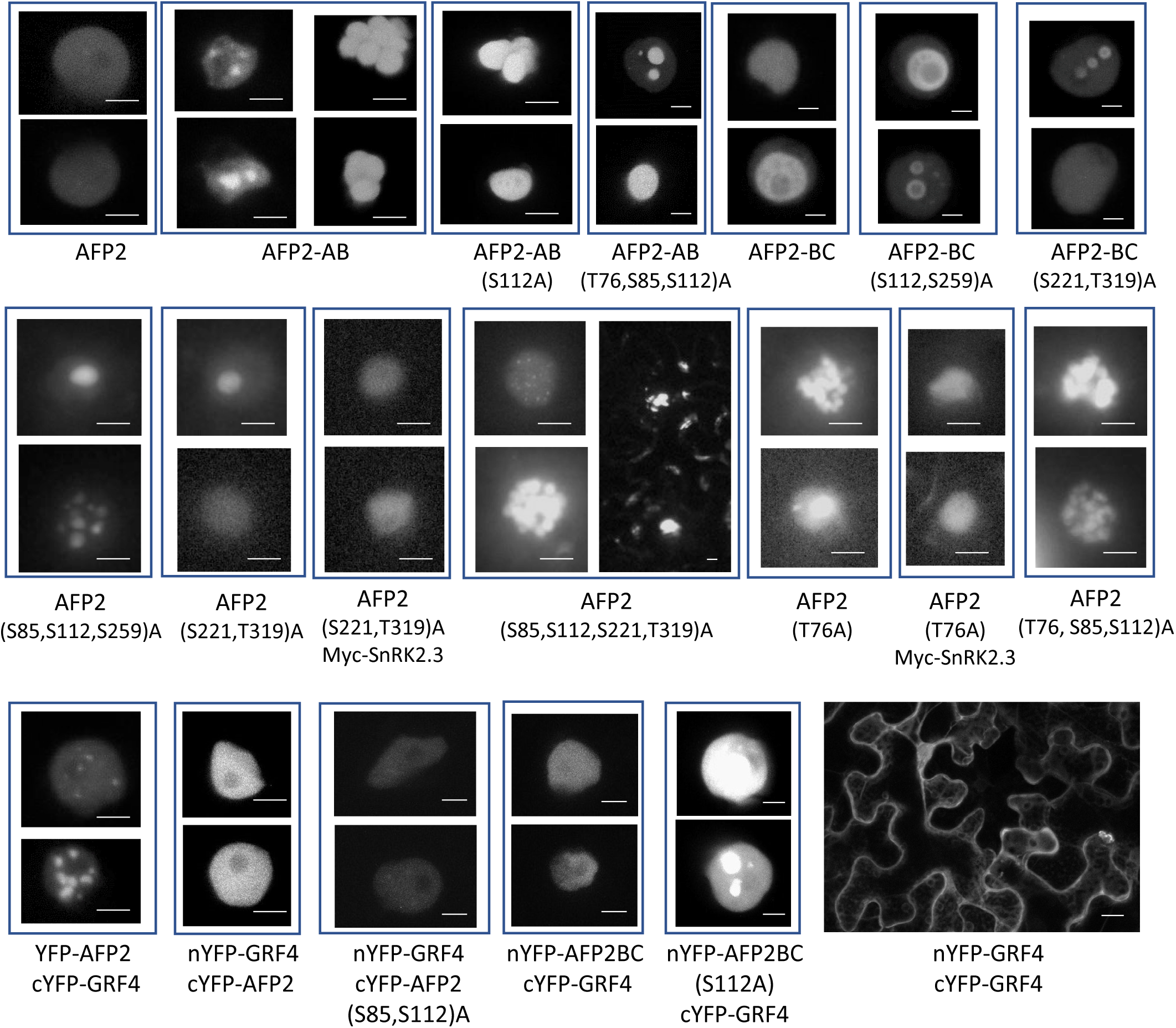
Localization of YFP-fusions to wild-type and phosphomutant AFP2-AB and –BC domains, and split YFP assays with GRF4, in transiently transformed *N. benthamiana*. Confocal images of nuclei: scale bars = 5 μm. nYFP-GRF4/cYFP-GRF4 image: scale bar = 20 μm

Full-length YFP-AFP2 fusions with ala substitutions in S85,S112, and S259 are generally diffuse in the nuclei, but occasionally form condensates. Similar to their effects on the BC domain, the full-length (S221,T319)A mutants are also diffuse with occasional foci within nuclei, but become more diffuse when coexpressed with SnRK2.3. Combining the (S85,S112,S221,T319)A mutations in a full-length protein results in many dispersed nuclear foci and some punctae at the cell periphery. Fusions with ala substitutions in T76 alone are sufficient to produce multiple nuclear foci, but become diffuse when co-expressed with SnRK2.3. Combinations including T76A and ala substitutions in SnRK2 targets produce many nuclear foci, whether or not SnRK2.3 is co-expressed.

Complexes of signaling proteins may be scaffolded by 14-3-3 proteins such as the GRFs, which recognize phosphorylation in several consensus motifs. The S85 and S112 residues of AFP2 are both SnRK2 target sites and predicted binding sites for 14-3-3 proteins. Similar to the yeast two-hybrid result, comparison of BiFC assays with GRF4 and either wild type AFP2 or a (S85,S112)A mutant show greatly reduced interactions with the mutant (Fig. 8). However, GRF4 still interacts with the S112A mutant of the AFP2-BC domain, consistent with the potential for additional 14-3-3 binding sites in this domain. Unlike the nuclear interactions with AFP2, GRF4 dimers are primarily cytoplasmic.

### Effects of AFP2 phosphorylation on ABA sensitivity of germination

We previously showed that overexpression of YFP fusions to either full-length AFPs or just the BC domains were sufficient to confer resistance to ABA inhibition of germination in transgenic Arabidopsis, but AB-domain fusions had almost no effect (Lynch et al., 2017). The full-length AFP2 fusions conferred up to 100-fold reductions in ABA sensitivity and produced desiccation intolerant seeds, while the AFP2-BC domain fusions conferred up to 30-fold reductions in ABA sensitivity and did not impair seed viability. Arabidopsis carrying transgenes with mutations either blocking phosphorylation or creating phosphomimetics in the SnRK2 target sites were tested for effects on ABA sensitivity of germination.

Although overexpression of full-length AFP2 with a variety of mutations permitted germination on up to 50 μM ABA, lines carrying phosphomimetic (S85,S112)E or non-phosphorylatable (S85,S112)A mutant YFP-AFP2 fusions differed from those with the wild-type fusion in that seedling greening was still greatly impaired by any ABA (Fig. 9AB and Supp Fig 16). Transgenic seeds overexpressing YFP-AFP2 with mutations in the C domain sites (S221,T319)A differed from those affecting the SnRK2 targets in that they were not able to germinate on media with ABA exceeding 3 uM, but greening was not impaired by this lower concentration of ABA (Fig. 9AB). In contrast, transgenic seeds overexpressing YFP-AFP2 combining all of these ala substitutions (S85,S112,S221,T319)A blocked seed maturation, resulting in production of green seeds lacking desiccation tolerance (Supp. Fig.17AB). ABA sensitivity of germination of seeds expressing this quadruple mutant protein was tested by removing maturing seeds from developing siliques. Although also able to germinate on at least 50 uM ABA, subsequent seedling development was severely impaired even in the absence of ABA (Supp. Fig.17CDE). The relatively weak ABA resistance of transgenic lines with the (S221,T319)A variant reflected the low level of fusion protein (Fig.7 and Supp. Fig. 17G). In contrast, the (S85,S112,S221,T319)A mutant constructs produced similar levels of fusion protein as the wild-type construct in seeds, and were highly effective in blocking ABA response, but the mutant protein was not maintained during seedling growth (Supp. Fig. 17EFG).

**Fig. 9.**
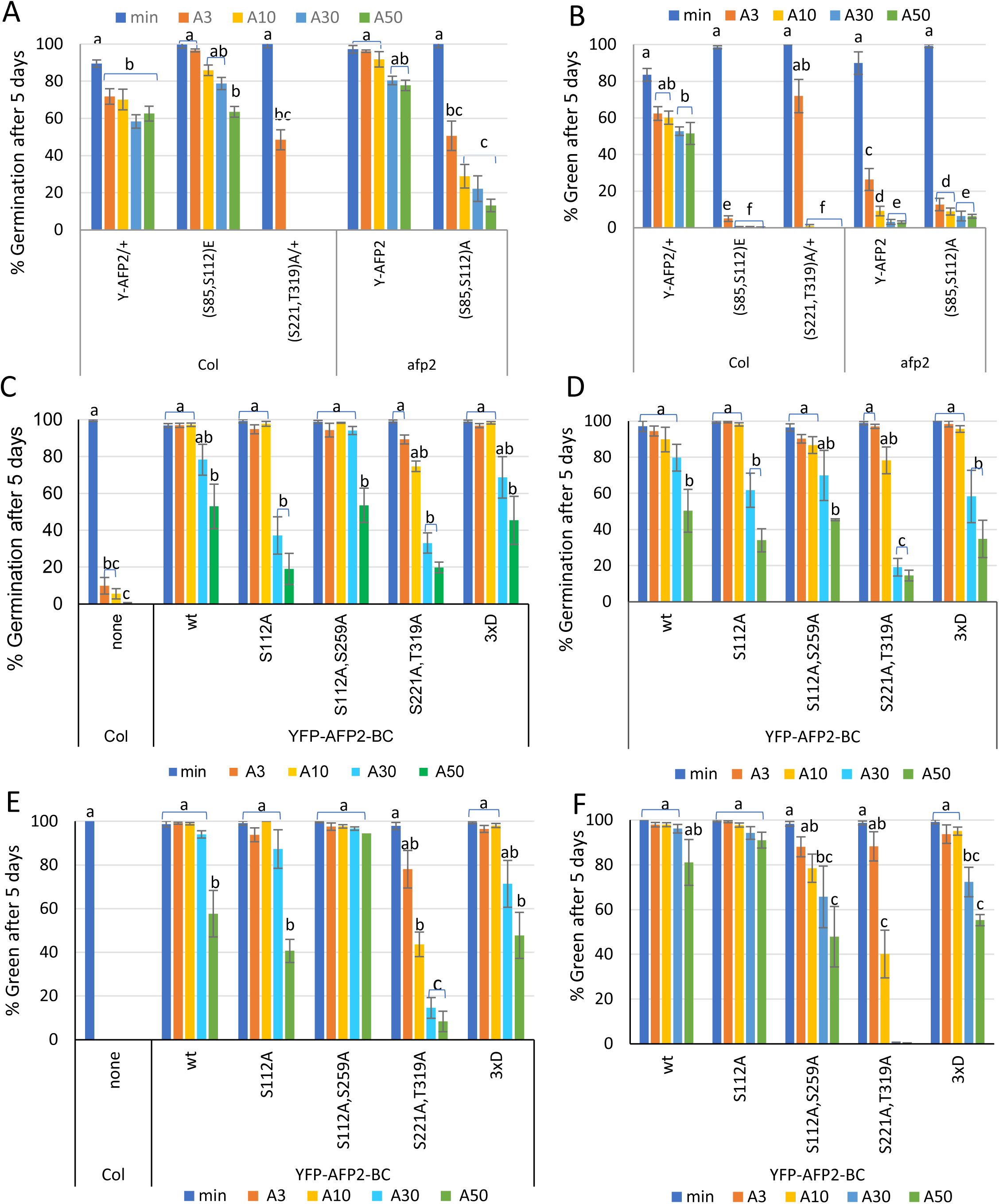
Physiological analysis comparing germination (A,C,D) or cotyledon greening (B,E,F) of transgenic lines overexpressing wild-type (wt) vs phosphomutant or phosphomimetic AFP2 fusions in wt(Col) background (A,B,C,E) or *afp2* background (A,B,D,F). Data displayed is the average of at least triplicate assays for each genotype and treatment ± S.E. Bars with different letters represent statistically different values using Tukey’s HSD post-hoc test (p<0.01).

The poor viability of seeds and seedlings expressing the full-length YFP-AFP2 resulted in partial silencing of these transgenes over generations, and led us to focus on AFP2-BC domain fusions for additional mutant analyses. When overexpressed in a wild-type background, all YFP-AFP2-BC domain fusions conferred similar resistance (Fig. 9C). The well-expressed transgenes from the wild-type background were backcrossed into the *afp2* mutant to create isogenic lines with similar transgene expression levels (Supp Fig. 13C), resulting in similar resistance to ABA as seen in the wild-type background up to 10 μM ABA (Fig. 9D). However, the mutants with alterations in the residues predicted to be phosphorylated by MPKs were less effective than the wild-type fusion in permitting germination on ABA higher than 30 μM in this background (Fig. 9D). Comparison of protein levels showed that only the (S221,T319)A mutant had reduced fusion protein accumulation, but accumulation could be increased by MG132 treatment (Fig.6E). Germination *sensu stricto* is defined as complete when the radicle emerges from the seed coat, and is usually followed by cotyledon greening during seedling establishment in the light. However, some ABA resistant lines including *abi5* mutants and AFP over-expressors often reverse this order when germinating in the presence of ABA, such that their cotyledons turn green prior to radicle emergence. Consequently, at the higher ABA concentrations, the fraction of seeds that have turned green may be higher than the fraction that has completed germination (Fig. 9EF). This was especially true for seeds expressing the wild-type and S112A mutants of the YFP-AFP2-BC fusion in the *afp2* mutant background. In contrast, the (S221,T319)A mutant was less effective than the other constructs in reducing ABA sensitivity in the *afp2* background and severely delayed greening when germinating on ABA.

### Effects of AFP2 phosphorylation on seed maturation

In addition to decreasing ABA sensitivity at germination, AFP2 overexpression inhibits various aspects of seed maturation including accumulation of storage proteins (e.g. cruciferins) and some late embryogenesis abundant (LEA) proteins (Lynch et al., 2022). To determine whether the phosphorylation state of AFP2 was important for this regulation, we compared storage protein accumulation in wild-type vs transgenic seeds with either wild-type or mutant AFP2 fusions. Surprisingly, even though most mutants were less effective than wild-type AFP2 for conferring ABA resistance, only the mutant forms with substitutions in multiple target residues resulted in a decrease in storage protein accumulation (Fig. 10, Supp. Fig.17H). The most dramatic reductions in cruciferin accumulation were seen in seeds producing the AFP2 quadruple mutant (S85,S112,S221,T319)A (Supp. Fig. 17H).

**Fig. 10.**
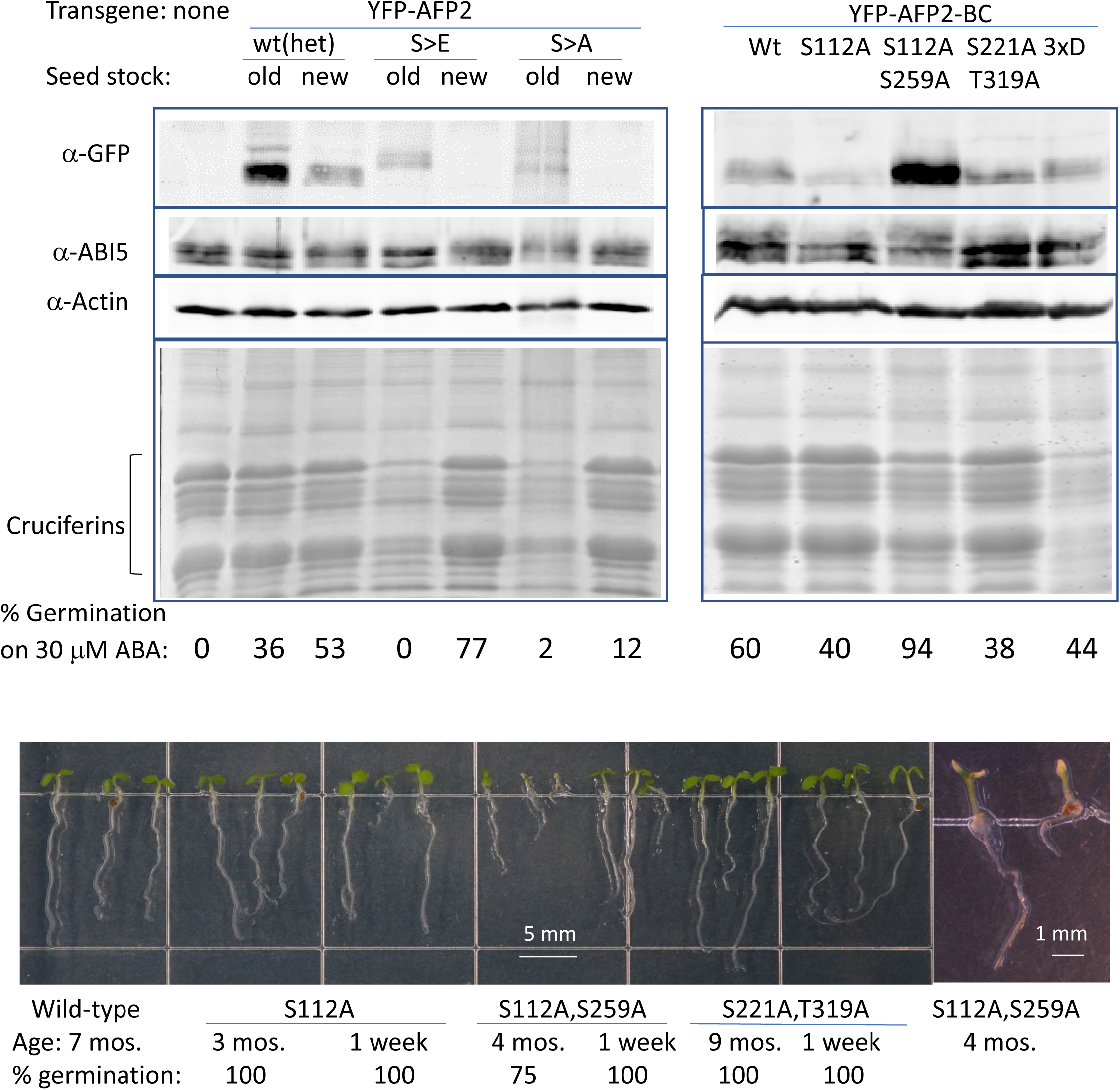
Effects of phosphomutant or phosphomimetic YFP-AFP2 fusions on storage protein accumulation and seed longevity. Top: comparison of dry seed protein extracts from lines either segregating a full-length wild type AFP2 fusion or homozygous for S85E,S112E (S>E) or S85A,S112A (S>A) mutant full-length AFP2 fusions (left) or AFP2-BC fusions with the indicated mutations (right). Upper panels show immunoblots of YFP-AFP2(-BC) fusions detected by α-GFP, -ABI5, and -Actin as a loading control. Lower panels show Coomassie staining of a comparably loaded higher 15% SDS-PAGE to retain and resolve the storage proteins. The “old” extracts were from early generations of these lines with high transgene expression and poor seed viability, while “new” extracts were from later generations with partially silenced transgenes permitting better viability but still conferring ABA-resistant germination. Bottom: seedlings from seed stocks of the indicated ages and genotypes, and their % germination, after 5d growth on minimal medium.

**Figure 11.**
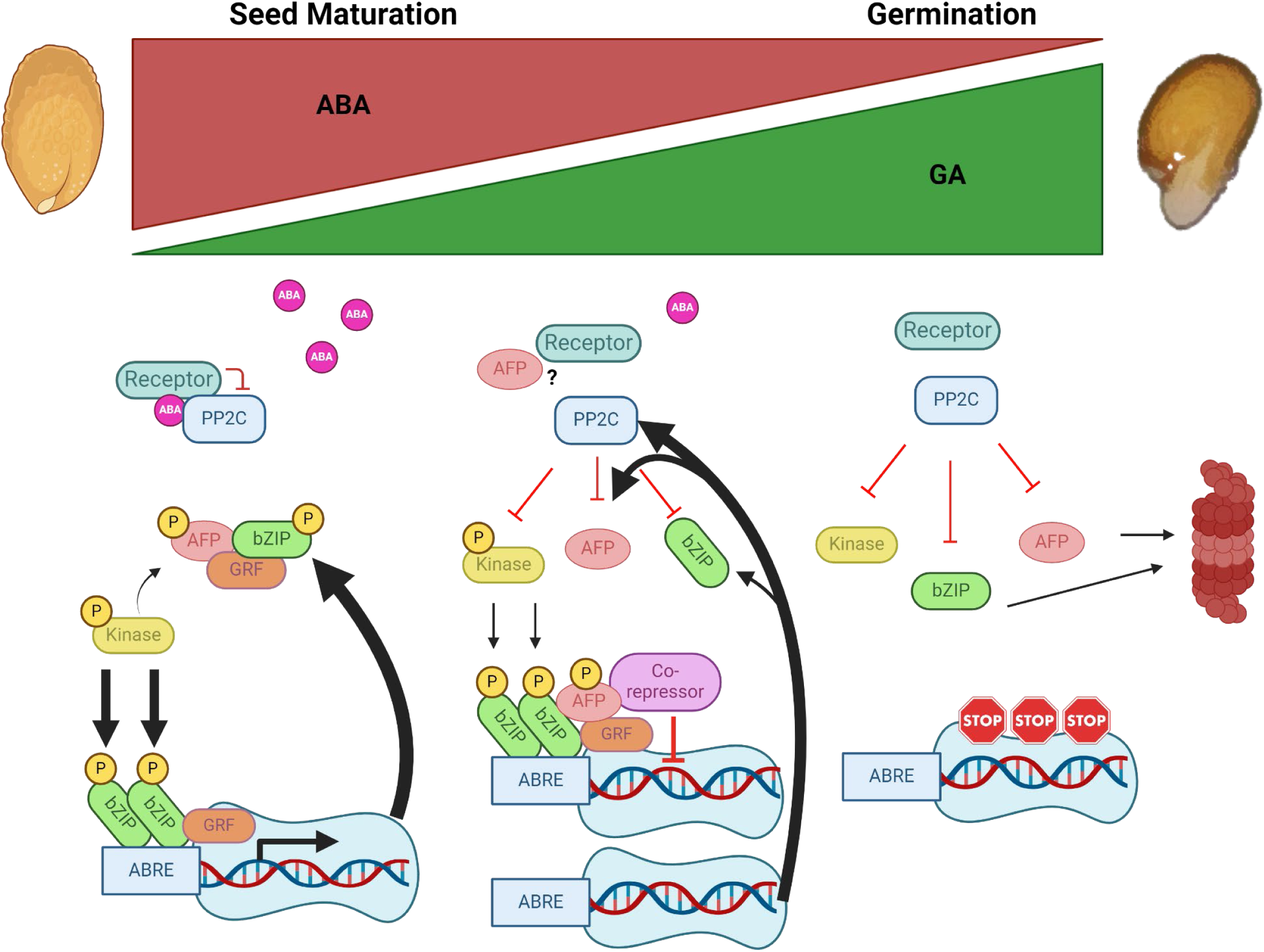
Changing relationships in the ABA core signaling pathway during the transition from seed maturation to germination. At high ABA concentrations, PP2Cs (e.g. AHG1 & AHG3) are inactive, kinases (e.g. SnRK2s & MPKs) are active, bZIPs (e.g. ABI5 & ABFs) and AFPs are phosphorylated and are bound by GRFs (14-3-3 proteins). When the bZIP:AFP ratio is high, ABA-regulated gene expression proceeds and germination is blocked (Garcia et al, 2008). As ABA levels decrease, ABA induces repressors including AFPs and PP2Cs in a feedback mechanism, the bZIP:AFP ratio decreases and seed-expressed gene expression declines. Interaction between AFPs and co-repressors (e.g. TPL/TPR and HDAC components) leads to inactivation of seed-expressed genes (Lynch et al. 2016). At low ABA levels, the PP2Cs are active and inactivate the SnRK2s, bZIPs (Lynch et al. 2012) and AFPs, and the latter two classes are proteasomally degraded. Many additional interactors are not included due to space constraints. The figure was created with BioRender.com.

The two BC domain mutant fusions producing the greatest reduction of storage protein accumulation ((S112,S259)A and “3xD”) had similar effects on ABA sensitivity of germination and seedling establishment (Figs. 9,10 and Supp Fig 16). However, they differ in that the (S112,S259)A mutations also adversely affect seed longevity, resulting in decreased viability within 4 months and complete lethality by 6 months (Fig. 10, Supp. Fig 18). It is noteworthy that the (S112,S259)A mutant form of AFP2-BC accumulates to higher levels in seeds than the wild-type construct (Fig.10), suggesting that this form might have greater effects on disrupting seed maturation, but this difference fades away during seedling growth (Supp Fig 19).

Examination of multiple ages of seed lots producing different AFP2-BC fusion variants showed that the major differences in storage protein levels reflected loss during aging (Supp Fig 18). Furthermore, this was not directly related to the level of ABI5 accumulation in that lines with very little storage protein still had ample ABI5. ABI5 is activated by phosphorylation and already appears as a triplet in dry seeds of all genotypes, but shifts to a more highly phosphorylated state when incubated on ABA (Supp Fig 19).

## DISCUSSION

### Post-translational modification of ABA core signaling components

The core signalosomes of ABA signaling comprise the PYR/PYL/RCAR receptors, clade A PP2C protein phosphatases, and SnRK2 kinases (reviewed in Lim et al., 2022; Née and Krüger, 2023). Each of these components is encoded by a multigene family that is differentially expressed, with different interaction characteristics and ABA sensitivities, creating the potential for a range of responses at different ABA concentrations in different tissues (Tischer et al., 2017). In the canonical view, ABA bridges the receptors with PP2Cs, effectively inactivating the PP2Cs and releasing the SnRK2s to become phosphorylated and active. When active, the SnRK2 kinases regulate the activity of many effector proteins, including the ABI5/ABF/AREB clade of bZIP transcription factors and ion channels. However, AHG1 differs from the other clade A PP2Cs in that it does not bind to the active sites of group 3 SnRK2s, interacts with only a subset of receptors and requires high ABA concentrations to do so (Tischer et al., 2017; Krüger et al., 2024). In addition to this central de-repression mechanism for activating ABA response by phosphorylation of effectors, all signalosome components are also subject to post-translational modifications including phosphorylation and ubiquitination. Both of these modifications may be either activating or inhibitory, depending on the target protein residues and the specific kinases or E3 ligases involved (reviewed in Lim et al., 2022).

Activity and stability of the ABI5/ABF/ABRE clade of bZIP transcription factors is also regulated by a variety of post-translational modifications including phosphorylation, ubiquitination, sumoylation, and S-nitrosylation (reviewed in Yu et al., 2015). Multiple kinases, including SnRK2s, CBL-interacting protein kinase (CIPK)26, SOS2-like protein kinase 5 (PKS5), BRASSINOSTEROID INSENSITIVE(BIN)2, and MPK3, phosphorylate overlapping and discrete subsets of amino acid residues within ABI5 (Wei et al., 2022; Lyzenga et al., 2013; Zhou et al., 2015; Bhagat et al., 2021). Our immunoblots detected multiple forms of ABI5 in dry seeds and seeds germinating in the presence of ABA, but it is currently not clear which modifications are present on these ABI5 proteins.

The AFPs interact with multiple members of this clade of bZIPs (Garcia et al., 2008) and AFP1 was initially proposed to assist degradation of these bZIPs by promoting interactions with E3 ligases that ubiquitinate ABI5 (Lopez-Molina et al., 2003). Although these proposed E3 ligase interactions appear weak at most (Lynch et al., 2022), subsequent studies including the present one have expanded the list of AFP interactors to include proteins representing all elements of the ABA signalosome, some MPKs, several 14-3-3 proteins, the DELLA repressors of GA signaling (Finkelstein and Lynch, 2022), the flowering regulator CONSTANS (Chang et al., 2019), WRKY36 and AHG1 in dormancy regulation (Deng et al., 2023; Krüger et al., 2024), several chromatin modifying factors (Pauwels et al., 2010; Lynch et al., 2017), and the protein kinase SALT OVERLY SENSITIVE(SOS)2 in regulating germination under salt stress (Wang et al., 2024). This extensive list of interactions is consistent with a role for AFPs as signaling hubs in a variety of processes, but raises the question of how specificity is achieved. Some of the likely options are regulated expression affecting availability of these proteins in different organs or stages of growth, and changes in phosphorylation state or other post-translational modifications leading to changes in structure of the disordered regions.

### Kinases regulating ABA response target AFPs

The kinase families most highly associated with ABA or osmotic stress signaling are the SnRK2 kinases acting in the ABA core signalosome, SnRK3/CBL interacting kinases (CIPKs), MAP kinases and calcium-dependent protein kinases (CDPKs). Many proteins can be phosphorylated at multiple sites, by multiple kinases, and the different sites may be phosphorylated sequentially with initial modifications altering the efficiency of subsequent events (reviewed in Miller and Turk, 2018). For example, mammalian GSK3 kinases often require “priming” by phosphorylation at a residue 4 or 5 amino acids C-terminal to the target (Sutherland, 2011). Furthermore, modifications at different sites may have opposing effects on activity of the substrate.

We focused on specific SnRK2s and MAP kinases that interacted with AFP1 and/or AFP2 in yeast two-hybrid and/or BiFC assays, using *in vitro* kinase assays and co-expression to test for altered mobility on SDS-PAGE and Phos-tag gels. Mass spectrometric analysis identified Ser-112 of AFP2 as the sole residue phosphorylated by SnRK2.6, consistent with results reported for a phosphorylated site conserved in all four AFPs (Krüger et al., 2024). Mutagenesis converting this residue to either alanine or a phosphomimetic amino acid reduced, but did not eliminate, phosphorylation of the AFP2-AB domain. This indicated that SnRK2.6 could phosphorylate additional residues, albeit not efficiently enough to be detected by mass spectrometry. Mutagenesis converting an additional predicted SnRK2 target, Ser-85, to alanine eliminated phosphorylation of the AFP2-AB domain. In contrast, combining the S112A mutation with alanine substitution of Ser-259, the residue with the next highest match to the SnRK2 consensus target in the AFP2-BC domain, was not sufficient to eliminate phosphorylation of this domain *in vitro*, but did block ABA-induced changes in mobility of the YFP-AFP2-BC fusion in stable transgenic plants. Mutations affecting predicted MAP kinase target residues did not block ABA-induced changes in mobility of protein fusions, but severely reduced stability and consequently accumulation of these proteins. Surprisingly, transient expression of full-length AFP2 fusions containing alanine substitutions at multiple documented or predicted kinase targets produced even more distinct phosphorylated products than the wild-type fusion. These mutant fusion proteins in stable transgenics were barely detectable, but some were slightly stabilized by inhibiting proteasomal degradation.

### AFPs can form biomolecular condensates

Predictions of AFP structures show a series of likely protein binding regions separated by intrinsically disordered regions (Lynch et al., 2017) and the AFPs have been reported to interact with diverse proteins. The structure predicted by AlphaFold for the AFP2 monomer shows most of the predicted SnRK2 and MAPK phosphorylation sites in regions predicted to be disordered (Supp. Fig.1). Only T319 is predicted with high confidence to be at the junction between beta-sheet and helical regions, while T76 and S112 are predicted to be near regions that might form helices. Our interaction data show that AFP2 forms dimers requiring regions in both the AB domain and the full-length protein (Supp. Fig.7). When modeled as a dimer, the relatively structured regions of the B domain (aa 115-143) and the C domain (aa 281-326) of distinct chains are in proximity, and T76 is predicted to be in a fully unstructured region. The AB domain does not dimerize and is predicted to have all potential phosphorylation sites in disordered regions. Substrates are often bound to the kinase active site in an extended conformation (reviewed in (Miller and Turk, 2018) and these structural predictions are consistent with the observation that the AB domain is more accessible to SnRK2s than full-length AFP2.

Intrinsically disordered regions are recognized as drivers of biomolecular condensate formation, often through multivalent interactions, i.e. with multiple other molecules (Emenecker et al., 2021). Post-translational modifications such as phosphorylation alter the charge distribution, and consequently the potential for interactions leading to condensate formation. Modeling possible effects of modification at the potential target residues tested in our mutant constructs showed limited impact of phosphorylation at either the two primary SnRK2 targets (S85 and S112) or the predicted MPK targets (S221 and T319), but additional phosphorylation of T76 or S259 was predicted to promote helix formation across much of the unstructured regions (Supp. Fig. 20). Further phosphorylation at additional target residues is predicted to give the AFP2 dimer even more structure, albeit with low confidence. In contrast, the ala substitutions tested in this study are predicted to maintain the dimer in a largely disordered state, such that additional residues might be accessible to additional kinases.

Initial descriptions of the AFPs showed a variety of localization patterns, depending on the experimental conditions. Transient over-expression of AFP(1)::CFP in either onion epidermal cells or Arabidopsis seedlings treated with ABA produced diffuse nuclear localization, but co-bombardment with 35S::ABI5 resulted in co-localization in nuclear bodies (Lopez-Molina et al., 2003). Stable expression of *AFP2pro:AFP2:GFP* in an *afp2* mutant background was also primarily nuclear, but excluded from nucleoli, in ABA-or NaCl-treated seedlings (Garcia et al., 2008). These conditions enhanced accumulation of both AFP2 and ABI5, but did not result in restriction to nuclear bodies, consistent with diffuse nuclear localization of AFPs likely to be in a phosphorylated state due to ABA or stress exposure. However, within 7 hrs of removal from these stress treatments AFP2:GFP was no longer visible in the nuclei, but still detected by immunoblot. In addition, some punctate fluorescence was visible in the cytoplasms of a few cells near the root meristem. Subsequent studies of interactions between AFPs and ABI5/ABF proteins by split YFP BiFC assays using transient over-expression in *N. benthamiana* also showed nuclear localization, but no foci (Lynch et al., 2017).

In the current BiFC assays, we found that co-expression of AFPs with kinases revealed interactions in nuclei, but excluded from nucleoli, whereas co-expression with PP2Cs resulted in formation of nuclear bodies over a broad range of sizes (Fig.4), suggesting that dephosphorylated AFPs were more prone to form condensates. However, YFP-AFP2 co-expressed with cYFP-GRF4 also formed numerous small nuclear condensates, consistent with interactions dependent on phosphorylation. In an effort to determine which residues were most important for regulating condensate formation, we tested localization of YFP-AFP fusions with alanine substitutions at known or predicted phosphorylation sites. Surprisingly, the wild-type YFP-AFP2-AB domain fusion formed clusters of nuclear bodies, similar to the “grapebunch-like” condensates observed for ARF proteins (Powers et al., 2019), but the alanine-substituted mutant AFP2-AB domain fusions formed fewer clusters. This might reflect formation of complexes by phosphorylation-dependent interactions with 14-3-3 proteins that are reduced in the mutants. In contrast, the wild-type YFP-AFP2-BC fusion was mostly diffuse, but YFP-AFP2-BC fusions with mutations preventing phosphorylation of the predicted kinase target residues were more likely to form condensates, sometimes with small internal areas of exclusion (donut-shaped). When co-expressed with the tagged PP2C phosphatase Myc-AHG1, full-length AFP2 formed punctae whether or not the SnRK2 or GRF target residues could be phosphorylated. The AFP2-AB domain also continued to form punctae when co-expressed with AHG1, but fewer punctae formed when AHG1 and the AFP2-BC domain were co-expressed. In combination with the previously described diffuse nuclear localization in ABA-treated roots, this suggests that condensate formation is negatively correlated with ABA-induced phosphorylation.

### Physiological significance of AFP phosphorylation state

The AFPs have recently been reported to function as a downstream element of DOG1 control of dormancy. Proteomic and genetic studies showed that DOG1 maintains dormancy by repressing AHG1 and AHG3 activity (Krüger et al., 2024). In non-dormant or *dog1* mutant seeds AHG1 dephosphorylates AFP2 at S112 and AFP1 at the corresponding conserved serine, allowing the AFPs to function as repressors of ABA response, resulting in germination. These results are consistent with our observation that ABA promotes a shift to the phosphorylated state, whereas co-expression with AHG1 has the opposite effect. Our studies used a gain of function approach, such that overexpression of the AFP domains and mutant forms leads to ABA resistance if the AFPs are physiologically active.

Functional tests of the physiological significance of these phosphorylation options focused on effects on ABA sensitivity of germination, cotyledon greening, storage protein accumulation, and seed longevity. We previously reported that overexpression of YFP-AFP2 could reduce ABA sensitivity nearly 100-fold and was so extreme in a wild-type background that homozygous transgenic lines failed to complete seed maturation and become desiccation tolerant (Lynch et al., 2017). Overexpression of the quadruple mutant (S85A,S112A,S221A,T319A), even though present at barely detectable levels, had a more extreme effect, also blocking maturation of heterozygous seeds. Homozygous YFP-AFP2 transgenes are tolerated in an *afp2* mutant background, where the total AFP2 concentration is lower, and YFP-AFP2-BC domain fusions are sufficient to reduce sensitivity only 30-to 50-fold, also resulting in viable seeds. The reduced impact of the truncated protein might reflect the loss of the EAR domain, and therefore interactions with TPL/TPR co-repressors. However, these fusions could still inhibit ABI5 and related AREB/ABF clade transcription factors via interactions with their C domains (Garcia et al., 2008). Further truncation of AFP2, removing the B domain, results in much weaker reduction of ABA inhibited germination (Lynch et al., 2017). Surprisingly, removal of the unstructured region between the B and C domains to produce the “LITTLE NINJA” microprotein creates a dominant negative effect on jasmonic acid signaling resulting in stunted growth, but this construct was not tested for effects on ABA sensitivity (Hong et al., 2020).

Comparison of endogenous AFP and ABI5 levels over a range of ABA concentrations showed that ABI5 is usually in excess post-stratification, ranging from 8-10 fold higher than AFP1/2 when exposed to 1 μM ABA, but increasing to as much as 30 fold higher after 5d post-stratification on media containing 10 μM ABA (Garcia et al., 2008). Extended incubation on the higher ABA concentrations led to accumulation of more AFP protein, effectively reducing the ABI5/AFP ratio, which was accompanied by increased germination. If this ratio is a sensor of physiological state that regulates germination, the severe disruption of the ratio by the AFP over-expression approach might be acting by titrating their interacting factors.

Mutations of the BC domain fusions converting only the SnRK2 targets, S112 or both S112 and S259, to alanine had minimal effect on ABA sensitivity of germination. However, the double mutant substantially reduced seed longevity. In contrast, mutants preventing phosphorylation of both predicted MPK target residues, S221 and T319, were less effective than the wild-type BC domain in conferring ABA resistance to concentrations higher than 10 μM ABA. This might reflect the reduced accumulation of this mutant form due at least in part to its reduced stability. Overexpression of phosphomimetic mutants did not confer greater ABA resistance than provided by the wild-type YFP-AFP2-BC. Although the mutants with the strongest effects on ABA sensitivity ((S112,S259)A and 3xD) had cruciferin levels similar to wild-type seeds when fresh, they showed accelerated loss of these storage proteins during shelf storage. Cruciferins have been found to serve as protectants from oxidative stress leading to seed deterioration during ageing (Nguyen et al., 2015), but the 3xD mutants remained viable despite a massive loss of cruciferins within a year, suggesting that they rely on alternative protectants. A more detailed comparison of the proteomes or metabolomes of these mutants might identify components critical for seed longevity.

Although phosphomimetic substitutions are often used to simulate phosphorylated forms of proteins, they frequently do not confer the expected activities, especially if the phosphorylation is a binding site for an adaptor protein, e.g. 14-3-3 proteins (Kozeleková et al., 2022; Dephoure et al., 2013). Consequently, it is not surprising that the alanine and aspartic acid substitutions have some similar effects since both types of mutations prevent true phosphorylation. Given that AFP2 phosphorylation is required to interact with the 14-3-3 proteins, this might represent a sequestered state induced by ABA, maintaining ABI5 and other downstream signaling factors in an active state. As ABA levels decrease post-stratification, increased activity of the AHG PP2Cs could release the AFPs from complexes with the 14-3-3 proteins, permitting more inhibitory interactions with ABI5/ABFs/AREBs.

Finally, what is the physiological significance of these condensates? In other well-characterized plant systems, condensates include miRNA processing bodies (Xie et al., 2021), light response complexes regulating translation (Jang et al., 2019), high temperature-induced de-repression of gene expression (Bohn et al., 2024), sensors of hydration regulating germination (Dorone et al., 2021), and cytoplasmic sequestering of inactive transcription factors (Powers et al., 2019). Thus, they may represent complexes required for activity or inactivity of the relevant proteins. In the current work, all AFPs or AFP domains tested were present as overexpressed fusions to all or part of YFP. A trivial explanation is that at least some of these simply reflect high protein concentrations produced by over-expression in transient systems. However, some fusions formed foci despite accumulating to barely detectable levels, different mutations or co-expression conditions led to different localizations, and the BiFC studies permit visualization of only the fusion pairs being tested. As discussed above, the AFPs interact with many different proteins and are likely to be modified at additional sites by kinases not yet tested. Consequently, it is possible that these AFPs, and their variants created by distinct post-translational modifications, are present in diverse complexes with different functions, i.e. acting as signaling hubs.

## Materials and Methods

### Plant Materials and Transgenes

Split YFP fusions for MPKs, GRFs and PYLs were constructed using the Gateway compatible pSITE-nEYFP-C1 (GenBank Acc# GU734651) and pSITE-cEYFP-C1 (Acc# GU734652) vectors and PCR products with attL ends added as described in (Fu et al., 2008), following manufacturer’s instructions for LR Clonase reactions (Invitrogen). The *35S::YFP:AFP2* fusion in pEarleyGate104 (Earley et al., 2006) and split YFP fusions for AFP1 (AT1G69260) and AFP2 (AT1G13740) were also constructed by LR Clonase recombinations, as described in (Lynch et al., 2017). The split YFP fusions for SnRK2s and AHGs were described in (Lynch et al., 2012). The *35S::YFP:AFP2* fusion lines in the wild-type (Col-0) and *afp2-1* (SALK_131676) (Alonso et al., 2003) mutant backgrounds were lines #4A2 and H1, respectively, reported in (Lynch et al., 2017). All genes included in this study are listed in Supp. Table 1. Mutations affecting potential phosphorylation sites were created by PCR using the high-fidelity polymerase ExTaq (TaKaRa Bio) and primers shown in Supp. Table 2, designed using PrimerX (https://www.bioinformatics.org/primerx/documentation.html), with plasmids encoding cDNAs for full-length AFP2 or specific subdomains as templates. The AFP2-AB domain includes aa 1-149 and the AFP2-BC domain includes aa 94-348. For multi-site mutations, fragments overlapping at the mutation sites were amplified, then annealed and amplified by attL primers at the end points to create Gateway-compatible fragments.

Arabidopsis plants were grown in pots in growth chambers under continuous light at 22°C. *Agrobacterium tumefaciens*-mediated direct transformation of constructs with mutations in potentially phosphorylated residues was performed by the floral dip method (Clough and Bent, 1998), followed by selection of BASTA-resistant seedlings on Germination Medium (GM: 0.5x MS salts and vitamins, 1% sucrose) supplemented with 8 μg/mL BASTA (“Finale”, AgrEvo Environmental Health). Homozygous lines were identified by production of 100% BASTA-resistant progeny. Following crosses between *35S:YFP:AFP2-BC* fusion lines and *afp2-1* mutants, YFP-AFP2-BC overexpressing progeny were selected by BASTA resistance and homozygous *afp2-1* segregants were identified by PCR-based genotyping, as described at http://signal.salk.edu/tdnaprimers.2.html.

### Yeast Two-Hybrid Constructs and Assays

Fusions between the GAL4 activation domain (AD) and full-length PP2C cDNAs were described in (Lynch et al., 2012). Fusions between the AD domain and full-length and partial AFP cDNAs were described in (Lynch et al., 2017). Fusions between the AD domain and 14-3-3 cDNAs were constructed by CRE-lox recombination between pUNI clones and the pACT2-lox vector (Liu et al., 1998). Fusions between the GAL4 DNA binding domain (BD) and either AFP or PYL cDNAs were constructed using the pGBKT7-DEST vector (Lu et al., 2010) and PCR products with attL ends. BD fusions with SnRK2.2, 2.3 and MPK6 were constructed by CRE-lox recombination using the pAS2lox vector and by LR Clonase reactions with the pGBKT7-DEST vector and pDONR clones for SnRK2.6, SnRK2.10, and MPK3.

BD fusions were transformed into yeast line PJ69-4A selecting for growth on yeast synthetic medium (YSM) without trp. AD fusions were transformed into Y187, selecting for growth on YSM without leu. Interactions were tested by matings between pairs of lines carrying BD-and AD-fusions; following overnight incubation on YPD plates, matings were replica plated to YSM lacking leu and trp to select for diploids or YSM also missing histidine (his) and, in some cases, adenine to screen for diploids that had activated the HIS3 and ADE2 reporter genes. Diploids carrying BD-PYLs and AD vector or AD-AFPs were grown in YSM lacking leu and trp, then diluted to an OD600 of 0.9, before making serial 10-fold dilutions that were spotted in replicates on YSM lacking lacking leu and trp, or also lacking his. Diploids combining BD-AFPs with AD vector or AD-GRFs were also analyzed by growth assays with serially diluted cultures.

### Plant growth conditions

Germination assays testing ABA sensitivity of age-matched seeds were performed on minimal nutrient media supplemented with ABA at concentrations over the range from 0-50 µM, as described in (Lynch et al., 2017). Accumulation of fusion proteins was assayed by immunoblots of seeds or seedlings harvested after 5d incubation on Germination Medium (GM: 0.5x MS salts and vitamins, 1% sucrose) or minimal media solidified with 0.7% agar, then subjected to ABA at the indicated concentrations, or 100 μM cycloheximide (CHX) (Acros Organics, #357420010) or MG132 (UBPBio, # F1101) for the indicated times.

For transient expression *Agrobacterium* lines expressing fusion proteins were combined with GV3101 expressing the P19 protein of tomato bushy stunt virus to enhance transient expression, as described in (Voinnet et al., 2003). Cultures of all bacteria to be used in infiltration were grown overnight in YEP media with appropriate selective antibiotics, diluted to an OD_600_ of 1.0 in 10 mM MgCl_2_, 10 mM MES pH 5.6, 0.2 mM acetosyringone (Aldrich #D134406) and rocked at room temperature for 2-3 hrs prior to mixing and infiltration of leaves on 2-3 week old *Nicotiana benthamiana* plants. For BiFC assays, Agrobacteria expressing cYFP-and nYFP-fusions were combined just prior to infiltration. Fluorescence was scored 2-4 days later, using an Olympus AX70 microscope or a Leica SP8 Resonant Scanning Confocal microscope. Confocal images were processed with Fiji (Image J) to produce maximum intensity Z-stack composites.

For stability assays of transiently transformed plants, infiltrated leaves were further infiltrated at 3d post-Agrobacterium infiltration with GM supplemented with 100 μM cycloheximide (CHX) or MG132, or a solvent control and incubated for 6 hrs prior to harvest.

### Immunoblots

Arabidopsis seedlings or *N. benthamiana* leaf tissue were ground directly in 2x Laemmli loading buffer (1.5-2 μL/mg FW), microfuged 10 min at 4°C to pellet debris, then the supernatants were boiled 5 min prior to fractionation by SDS-PAGE or SuperSep**™** Phos-tag**™** PAGE (Fujifilm Wako Chemicals). Proteins were transferred to nitrocellulose filters (Prometheus, Genesee Scientific, San Diego, CA) and stained with Ponceau S, as described in (Lynch et al., 2017). Filters were blocked with Casein blocking buffer (LI-COR Biosciences, Lincoln, NE), then co-incubated with anti-GFP mAb (1:10000, UBPBio, Aurora, CO) and/or anti-actin mAb (A0480, Sigma) primary antibodies, followed by anti-mouse secondary IRDye 800 conjugated IgGs (LiCor, #926-32210), and visualized using the 800 channel of the Licor Odyssey Infrared Imaging System. Mature dry Arabidopsis seeds were ground in 20 μL/mg FW 1x Laemmli loading buffer, then processed as above. ABI5 was detected with anti-ABI5pAb (1:10000, Ab98831, AbCam) primary antibodies, followed by anti-rabbit secondary IRDye 700 conjugated IgGs (LiCor #926-68071), and visualized using the 700 channel of the Licor Odyssey Infrared Imaging System.

### Recombinant protein purification

Fusions between His6-tags and full-length, partial and mutant AFP coding sequences were constructed by CRE-lox recombination between pUNI cDNAs and the pHB3-His6 vector, as described in http://signal.salk.edu/UPS_Protocols_Vectors.pdf. Primers used for subcloning and mutagenesis are shown in Supplementary Table 1. Plasmids carrying fusion genes were transformed into *E.coli* BL21C+ and expression was induced by growth overnight at 19-20oC in Overnight Express^TM^ Instant TB medium (Novagen). Cells were harvested by microcentrifugation for 3 min at 16K rpm, resuspended in 140 μL Lysis Buffer (8.3 M urea, 0.1 M NaPhosphate pH 8) per mL of initial culture. His6-fusions were affinity purified in batch mode with HIS-select Nickel Affinity beads (Millipore P6611) according to the manufacturer’s protocol for denaturing conditions, eluting with 8 M urea, 0.1 M NaPhosphate pH 5.5. The urea concentration was reduced ten-fold for use in kinase reactions.

GST fusions to full length SnRK2.6 also made use of the pUNI vector system, and induction overnight in TB Express, but proteins were extracted under native conditions with CelLytic B in 1x PBS, 10 mM DTT, 83 units/mL Benzonase nuclease, 1 mM PMSF, 5 mg/mL lysozyme, followed by purification on glutathione-agarose beads (UBP-Bio, P3060-10) and elution with 30 mM Tris pH9, 300 mM NaCl, 5 mM DTT, 0.5% Triton-X100, 1 mM PMSF, 40 mM glutathione.

### In vitro kinase assays

Purified GST-SnRK2.6 was activated by 1 hr pre-incubation in kinase buffer (25mM Tris pH 7.5, 12 mM MgCl_2_, 2 mM DTT) with 1 mM ATP, then combined with His-AFP fusions maintaining the buffer and ATP concentrations for 1 additional hr of incubation. Phosphorylation was monitored in wild-type and mutant constructs by incorporation of ^32^P from gamma-labeled ATP (0.1-0.2 μCi/rxn) during the latter incubation. SDS sample buffer was added to 1x final, samples were boiled 5 min., then fractionated on 13% SDS-PAGE, dried and autoradiographed. Following assays with only cold ATP, gel slices containing lower mobility products were excised and sent to UC Davis for mass spec analysis. Additional SnRK assays with mutant AFP fusion proteins were analyzed by ProQ Diamond Phosphoprotein staining according to manufacturer’s instructions (Invitrogen).

### Mass Spectrometry

Coomassie stained gel slices were processed with a standard washing, reduction/alkylation followed by tryptic digestion, as described in <https://www.promega.com/-/media/files/resources/protocols/technical-bulletins/101/proteasemax-surfactant-trypsin-enhancer.pdf> The mass spectrometer used was a Xevo G2 QTof coupled to a nanoAcquity UPLC system (Waters, Milford, MA). Samples were loaded onto a C18 Waters Trizaic nanotile of 85 um × 100 mm; 1.7 μm (Waters, Milford, MA). The column temperature was set to 45°C with a flow rate of 0.45 mL/min. The mobile phase consisted of A (water containing 0.1% formic acid) and B (acetonitrile containing 0.1% formic acid). A linear gradient elution program was used: 0–40 min, 3–40 % (B); 40-42 min, 40–85 % (B); 42-46 min, 85 % (B); 46-48 min, 85-3 % (B); 48-60 min, 3% (B).

Mass spectrometry data were recorded for 60 minutes for each run and controlled by MassLynx 4.2 SCN990 (Waters, Milford, MA). Acquisition mode was set to positive polarity under resolution mode. Mass range was set from 50 – 2000 Da. Capillary voltage was 3.5 kV, sampling cone at 25 V, and extraction cone at 2.5 V. Source temperature was held at 110C. Cone gas was set to 25 L/h, nano flow gas at 0.10 Bar, and desolvation gas at 1200 L/h. Leucine–enkephalin at 720 pmol/ul (Waters, Milford, MA) was used as the lock mass ion at m/z 556.2771 and introduced at 1 uL/min at 45 second intervals with a 3 scan average and mass window of +/- 0.5 Da. The MSe data were acquired using two scan functions corresponding to low energy for function 1 and high energy for function 2. Function 1 had collision energy at 6 V and function 2 had a collision energy ramp of 18 − 42 V.

Raw data was searched using Protein Lynx Global Server 3.0.3 Expression 2.0 (Waters, Milford, Massachusetts). Processing parameters consisted of a low energy threshold set at 200.0 counts, an elevated energy threshold set at 25.0 counts, and an intensity threshold set at 1500 counts. A dedicated database was constructed containing common housekeeping proteins as well as the provided sequence of the target protein. The database was randomized within PLGS. Possible structure modifications included for consideration were methionine oxidation, carbamidomethylation of cysteine, deamidation of asparagine or glutamine, dehydration of serine or threonine, and phosphorylation of serine, threonine, or tyrosine.

### qRT-PCR analysis

RNA was isolated from *N. benthamiana* leaf tissue or Arabidopsis seedlings as described in (Lynch et al., 2017). DNA-free total RNA was prepared by incubation with RQ1 DNase (Promega) and RNAsin for 15 min. at room temperature, followed by inactivation of the DNase by addition of EGTA and incubation for 5 min. at 65°C, then clean-up over Qiagen RNeasy or Zymo-Clean columns according to manufacturers’ instructions. Approximately 0.5 μg of RNA was used as template for cDNA synthesis by MMLV reverse transcriptase (Promega), using a 10:1 mix of random hexamers and oligo dT as primers. cDNA concentrations were checked by qRT-PCR, using Forget-Me-Not master mix (Biotium) in an iQ5 or CFX cycler (Bio-Rad), according to manufacturer’s instructions. Primers used for normalizing were selected for uniform expression in Arabidopsis seeds or young seedlings subjected to a variety of treatments: AT5G46630 and At1g13320 (Czechowski et al., 2005) or a validated reference gene for *N. benthamiana* (Pombo et al., 2019). Equal amounts of cDNA were used as templates for reactions with all other primer sets (listed in Supp. Table 1), quantified relative to a standard curve spanning the range of concentrations present in all genotypes of transgene expression.

## Acknowledgements

We thank the ABRC team at Ohio State University for efficient distribution of vectors and cDNA clones, Armann Andaya at the UC Davis Campus Mass Spectrometry Facilities for MS analysis, Ben Lopez at UCSB for confocal microscopy, and Delaney O’Donnell for assistance with screening transgenic lines. This work was supported by a UCSB Academic Senate Grant to RRF, Faculty Research Assistance Program funds, and National Science Foundation Grant# IOS1558011 to RRF.

## Author contributions

RRF conceived the project and supervised construction of genetic lines and physiological assays; all authors performed experiments; RRF wrote the article, with contributions from BJEM and TL, and serves as the author responsible for contact.

## Supplementary Figures

**Supp. Fig. 1.**
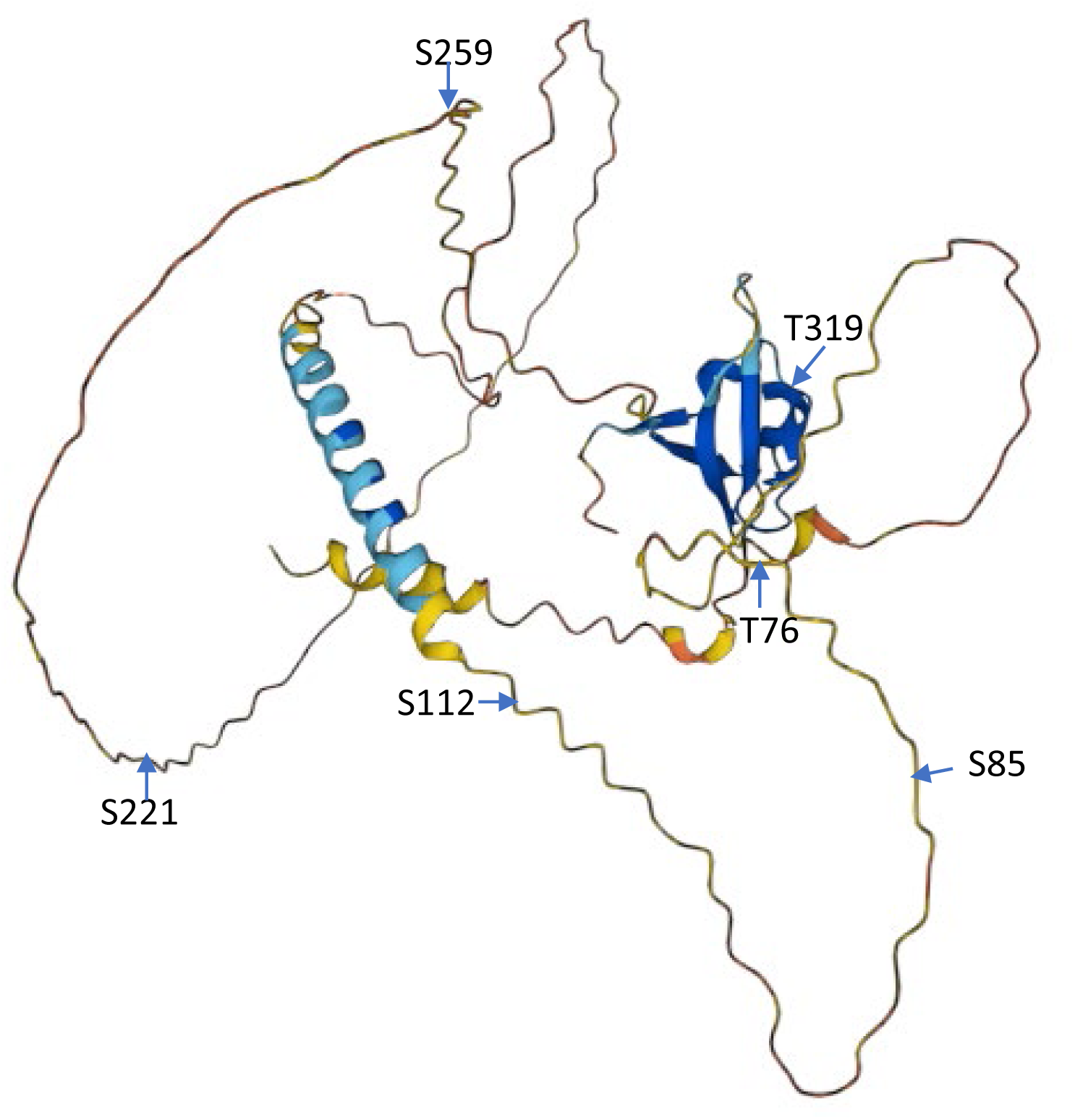
Alphafold prediction of AFP2 structure (Model AF-Q9LMX5-F1) with locations of predicted SnRK2 or MPK phosphorylation sites labeled.

**Supp Fig. 2.**
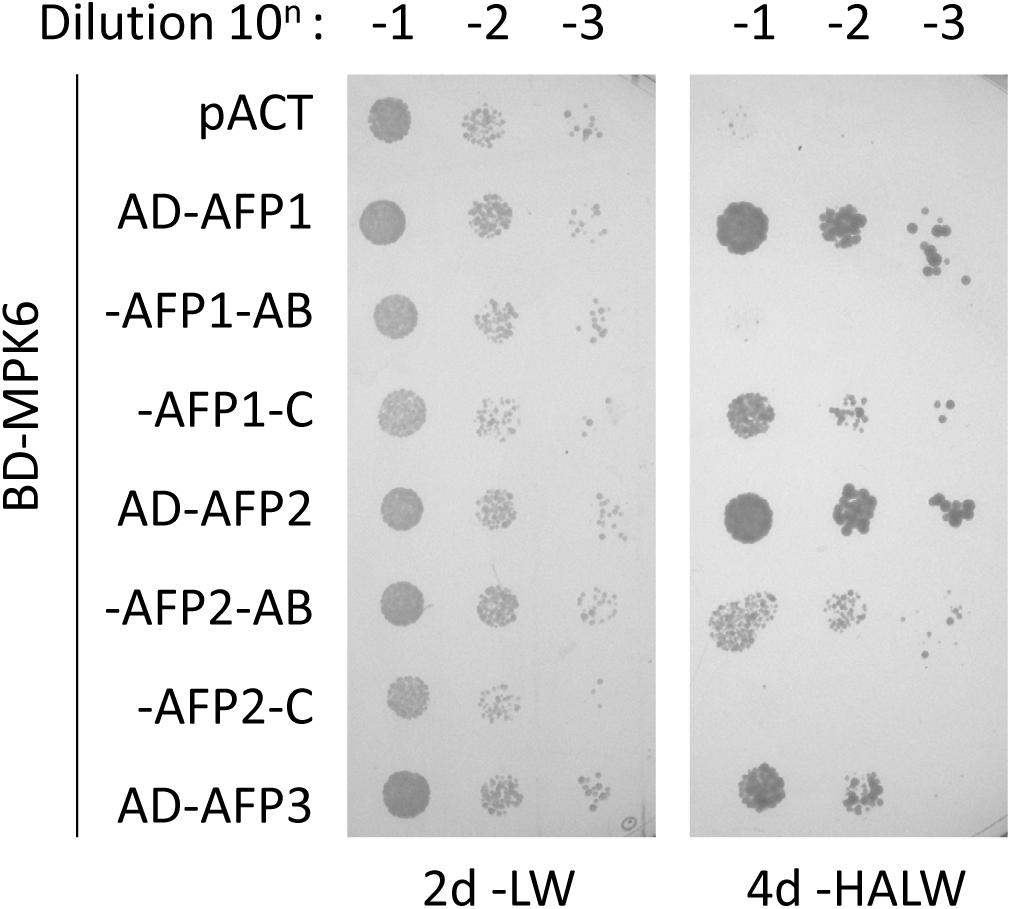
Interactions between MPK6, AFPs and their subdomains detected by yeast two-hybrid assays. Following matings between haploid lines carrying the BD-MPK6 or AD-AFP fusions, the indicated diploid lines were grown overnight in media lacking leu and trp. After measuring the OD600, cultures were diluted to the same concentration, then serially diluted 10-fold three times. The diluted cultures were then replica-spotted onto selective media. The -LW plate serves as a control for accuracy of the dilutions. Growth on the -HALW plate requires interaction between the AD- and BD-fusions to activate the reporter genes complementing the his3- and ade2-mutations of these yeast.

**Supp Fig. 3.**
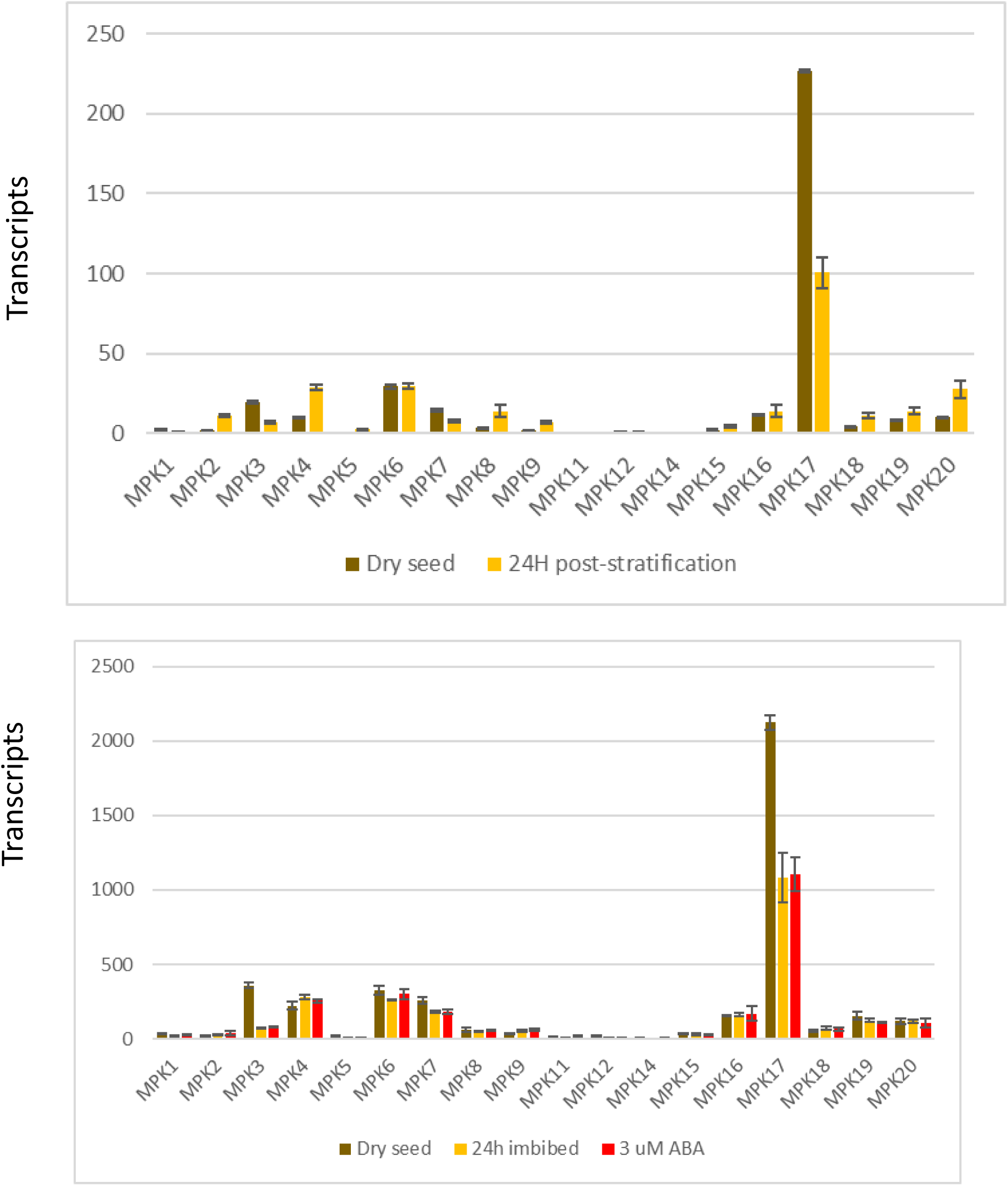
Expression of MPKs in dry seeds and either 24 h post-2d-stratification (top) or after 24h imbibition in water or 3 uM ABA following 2-4 months after-ripening (bottom). Data from (Narsai et al., 2011; Nakabayashi et al., 2005) displayed as “Absolute” units on http://bar.utoronto.ca/efp//cgi-bin/efpWeb.cgi (Winter et al., 2007).

**Supp. Fig. 4.**
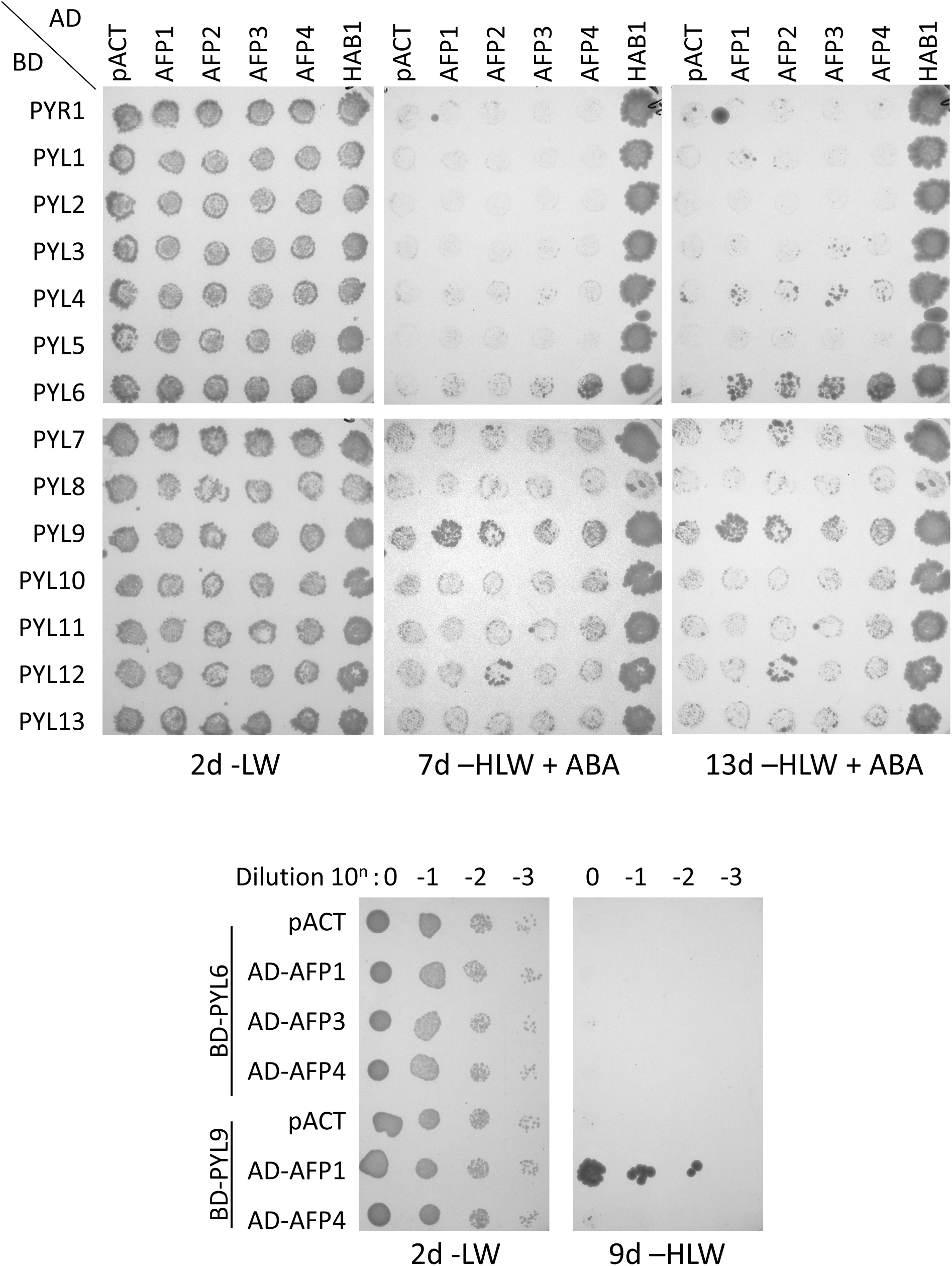
Yeast two-hybrid test of interactions between ABA receptors and AFPs, presented as GAL4BD-PYR/PYL and GAL4AD-AFP fusions. The AD fusion to the PP2C HAB1 is included as a positive control, and pACT is a negative control. Fusions were combined by matings between Y187 carrying AD fusions and PJ69-4a carrying BD fusions. After overnight incubation on YPD media, yeast were replica plated onto selective media lacking leu and trp (-LW) to maintain the AD-and BD-fusion plasmids, then replica plated again to media lacking histidine (-HLW) and supplemented with ABA to score interactions between the fusion proteins. Lower panel: test of potential positives by dilution assay, as in Fig.3.

**Supp. Fig. 5.**
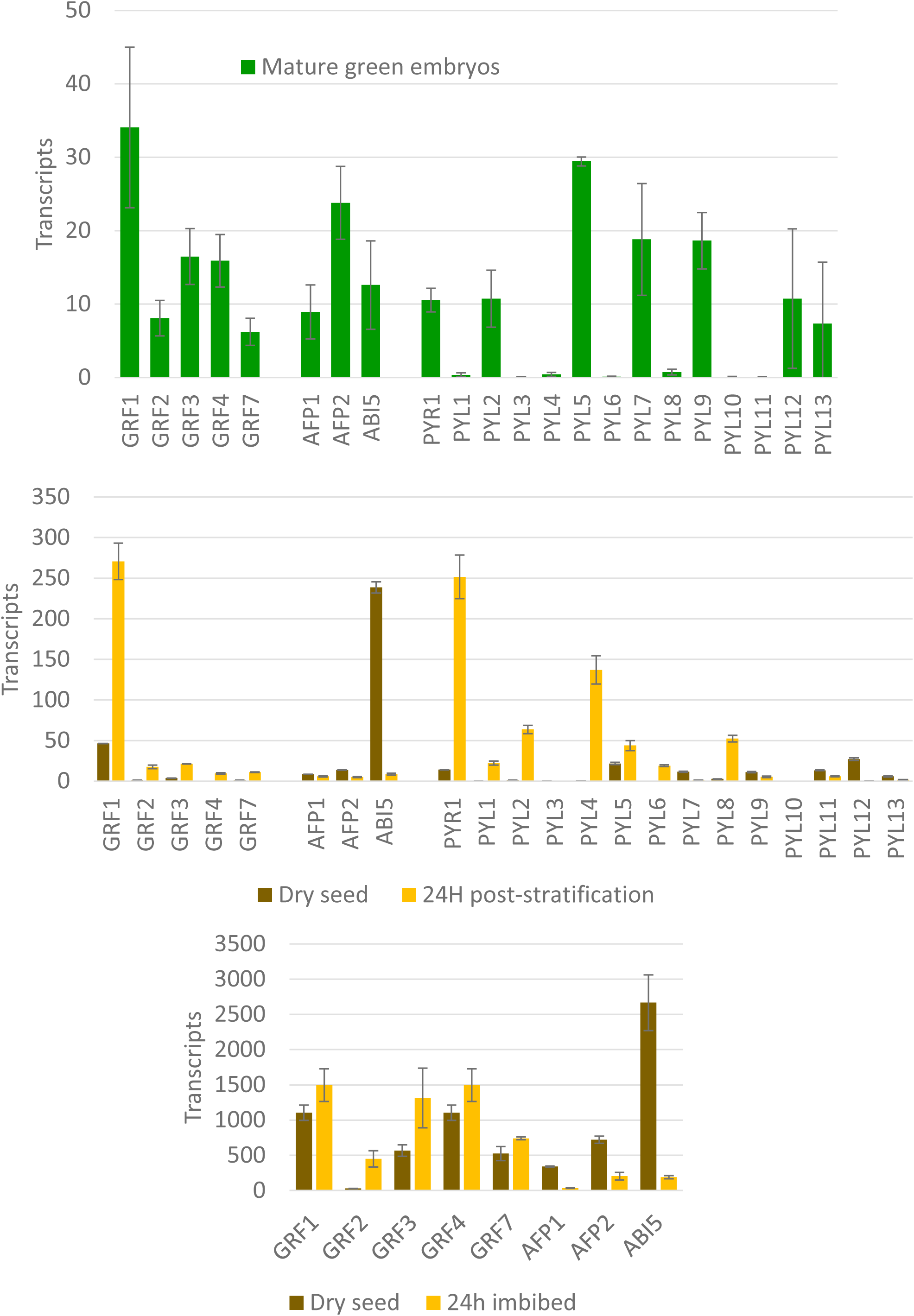
Expression of PYLs, GRFs and AFPs in maturing embryos (top), dry seeds and either 24 h post-2d-stratification (middle) or after 24h imbibition in water following 2-4 months after-ripening (bottom). Data from (Narsai et al., 2011; Nakabayashi et al., 2005; Hofmann et al., 2019) displayed as “Absolute” units on http://bar.utoronto.ca/efp//cgi-bin/efpWeb.cgi (Winter et al., 2007)

**Supp. Fig. 6.**
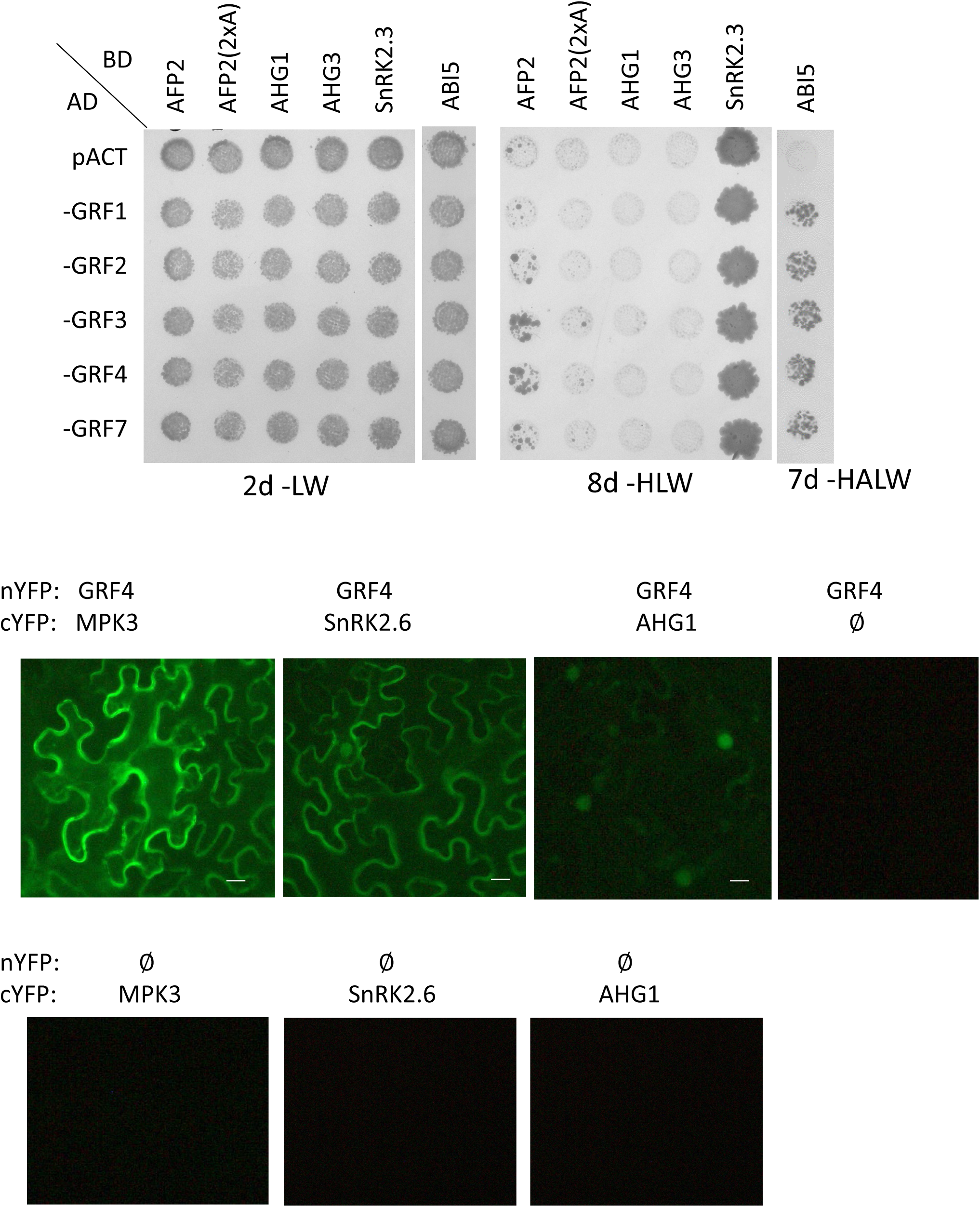
Interactions between 14-3-3 proteins (GRFs) and ABA core signaling elements. Top: Yeast two-hybrid assays combining AD-fusions to 14-3-3 proteins with BD-fusions to wild-type and mutant AFP2, PP2Cs, SnRK2.3 and ABI5 by matings, as described for Supp.Fig. 4. Bottom: BiFC assays testing GRF interactions with kinases and phosphatases. Scale bars = 10 μm

**Supp Fig. 7.**
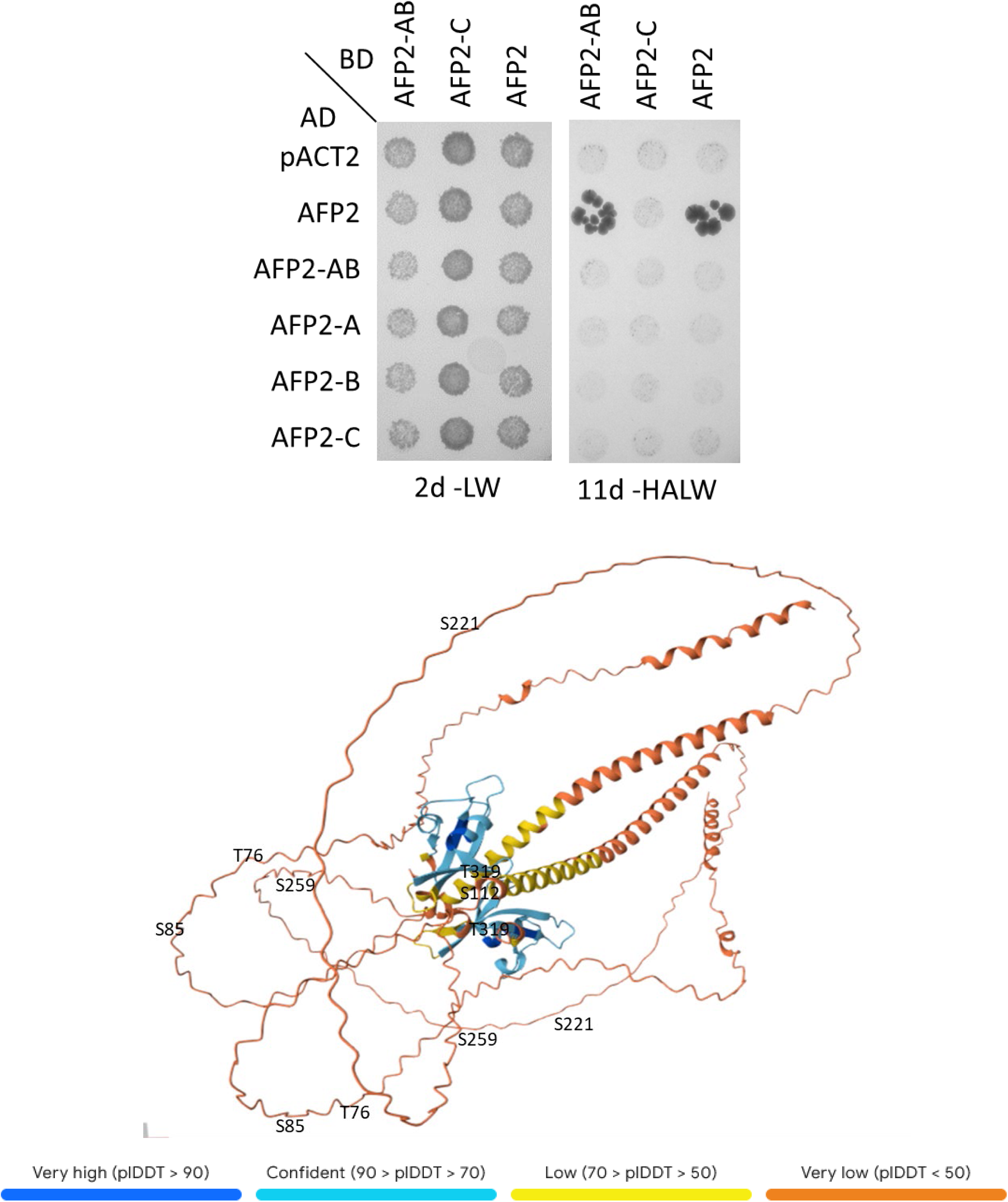
Potential dimerization domains of AFP2 Top: Mapping interacting domains of AFP2 by yeast two hybrid assays, as in Supp. Fig. 2. The AFP2-AB domain includes aa 1-149 and the C domain includes aa 150-348. Bottom: Alphafold prediction of AFP2 dimer structure.

**Supp. Fig. 8.**
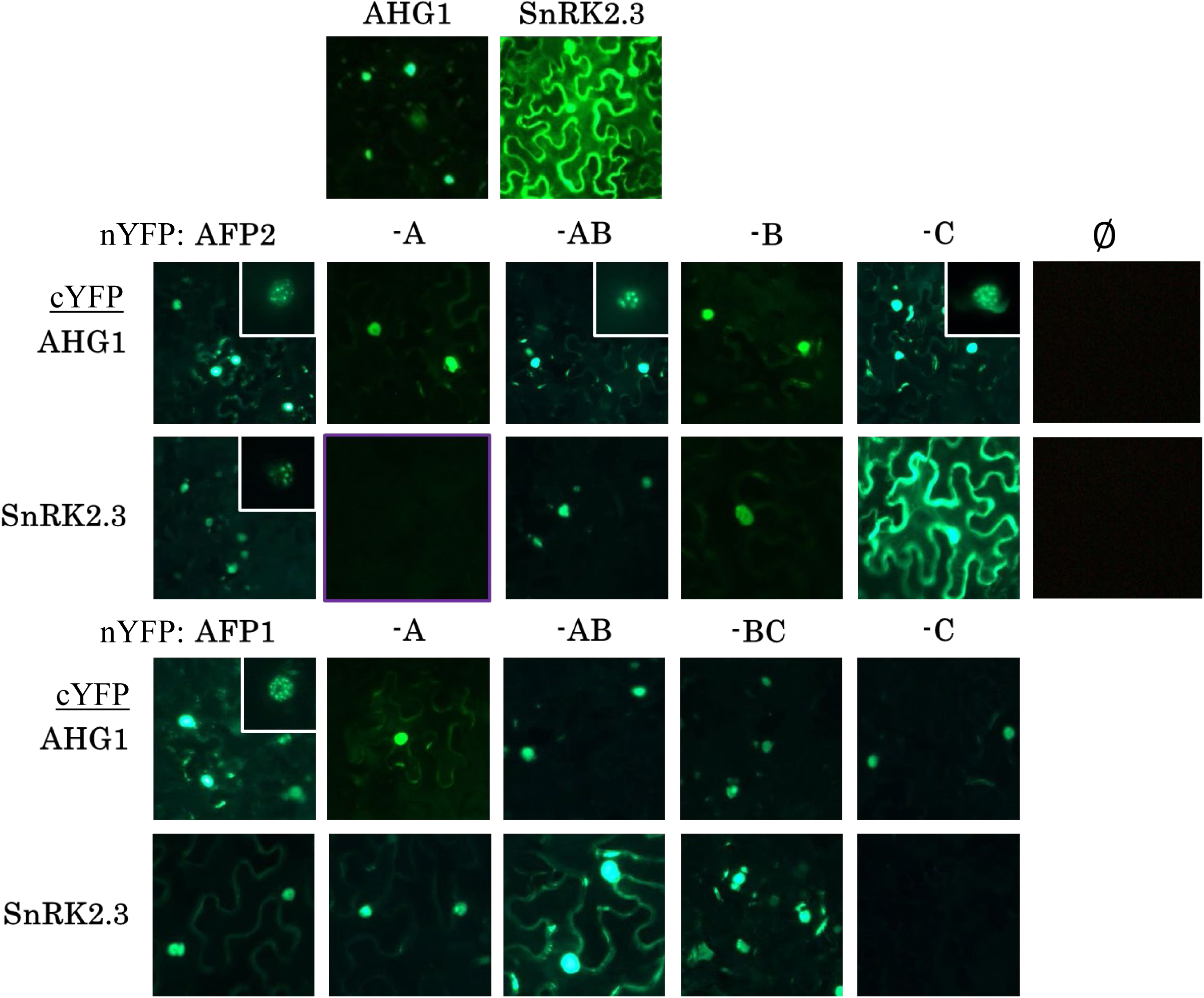
Biomolecular fluorescence complementation of full length AFP2 (middle) and AFP1 (bottom) and subdomains versus ABA core signaling components in *N. benthamiana* leaves (100X magnification). Top panel shows localization of YFP-fused phosphatase and kinase for reference. Color differences are due to different versions of image capture software. Inset images show examples of nuclear speckles observed at 200X or 400X magnification (Erickson McNally, 2016)

**Supp. Fig. 9.**
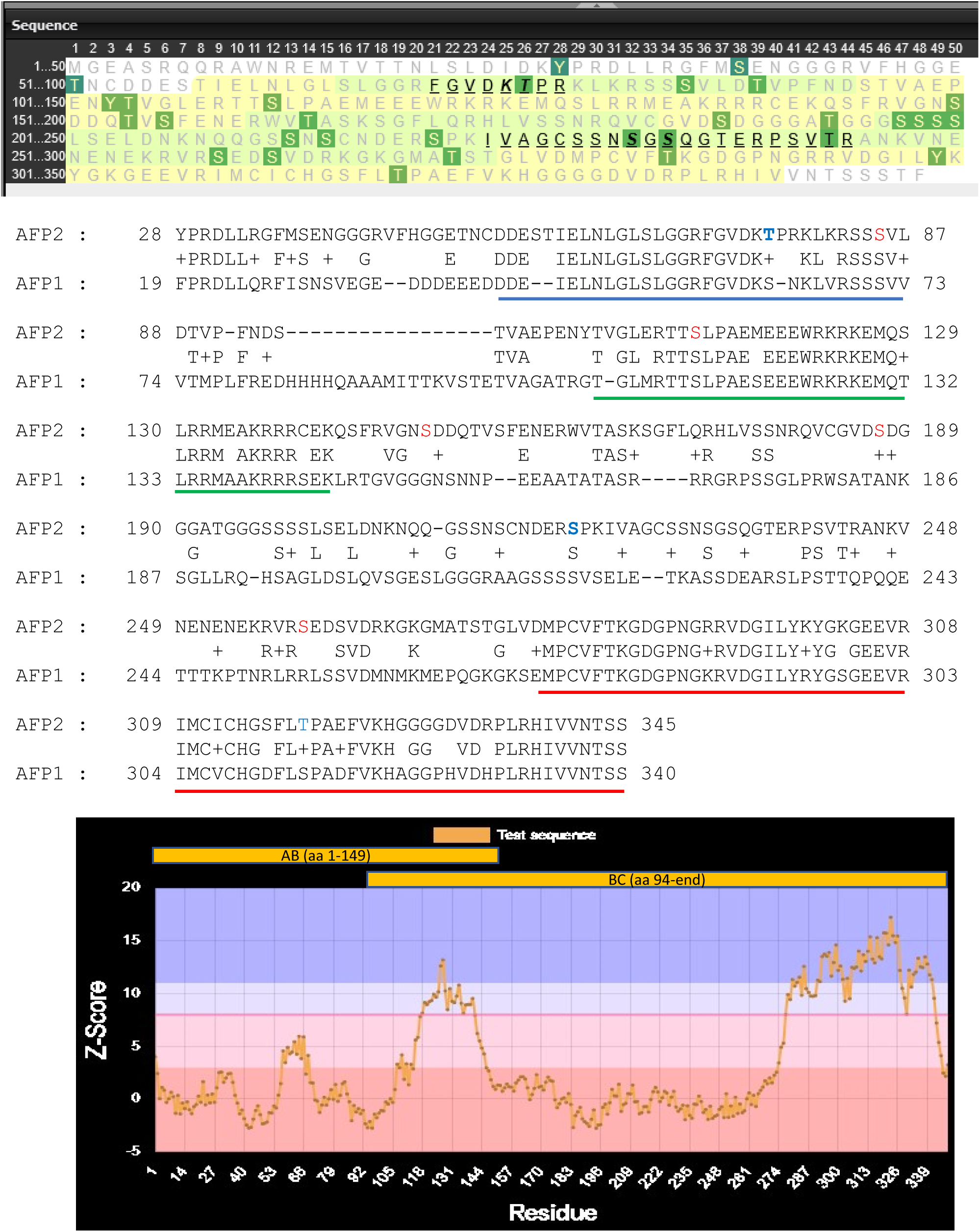
Predicted phosphorylated residues in AFP1 and AFP2. Top: PhosPhAt shows possible phosphorylation sites of AFP1 highlighted in green, with documented sites identified by black letters. Middle: Alignment of AFP1 and AFP2. A, B and C domains are underlined in blue, green and red, respectively. Residues fitting consensus for MAPK phosphorylation are in blue, those fitting SnRK2 consensus are in red. Bottom: Disordered regions (score <8) in AFP2 predicted by Attention based DisOrder PredicTor (ADOPT) (Redl et al., 2023).

**Supp. Fig. 10.**
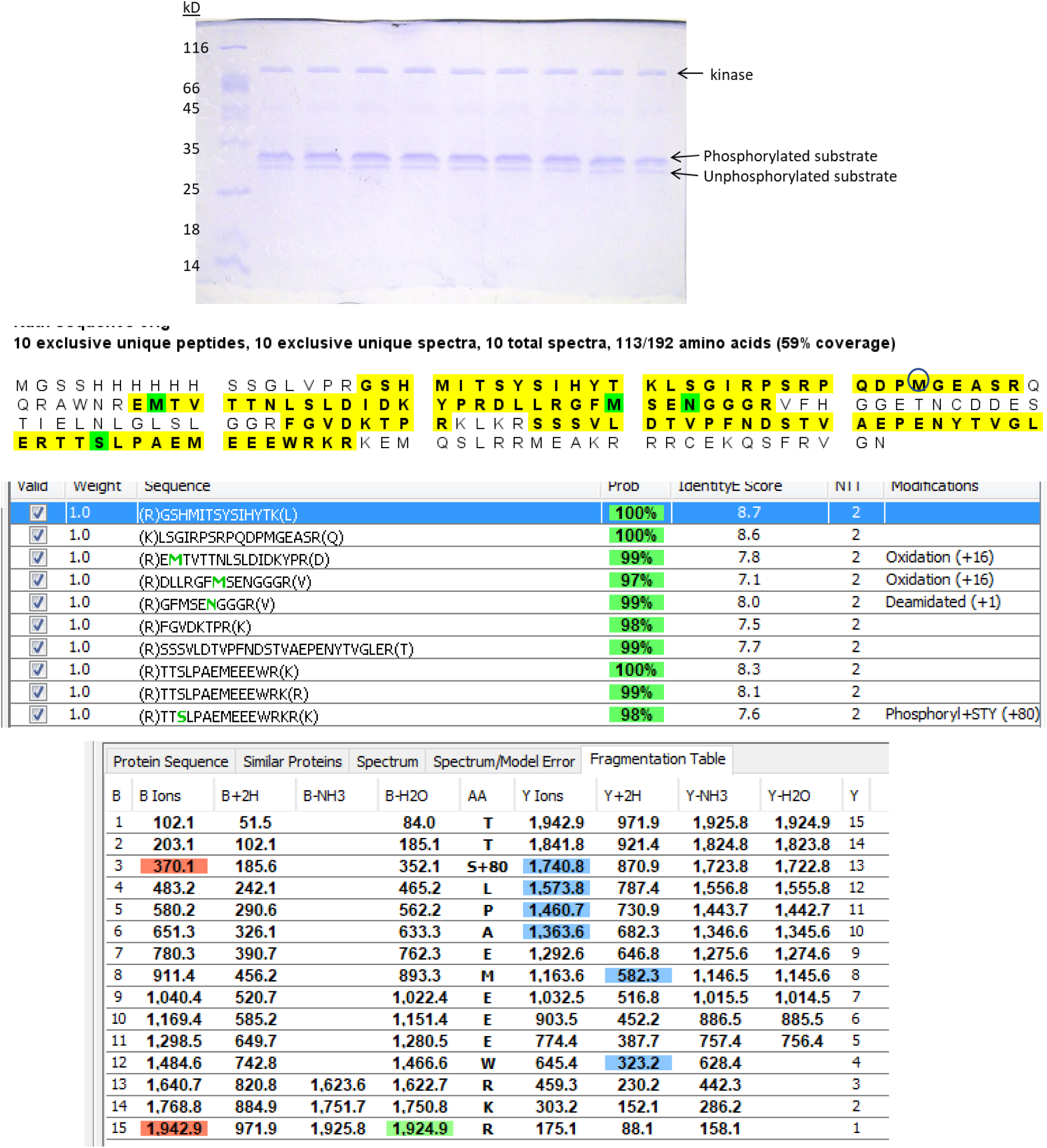
*In vitro* kinase reaction of His-AFP2-AB incubated with GST-SnRK2.6 used for mass spectrometric analysis of phosphorylation site(s). Top: Coomassie-stained SDS-PAGE; the replicate phosphorylated substrate bands were excised from the gel, and sent to the UC Davis Mass Spectrometry Facility for analysis following tryptic digestion. Bottom: Summary of Mass Spec analysis by Scaffold software, including Fragmentation Table for the only peptide showing phosphorylation. The initiating Met of AFP2, at position 44 of the fusion protein, is circled in the sequence.

**Supp. Fig. 11.**
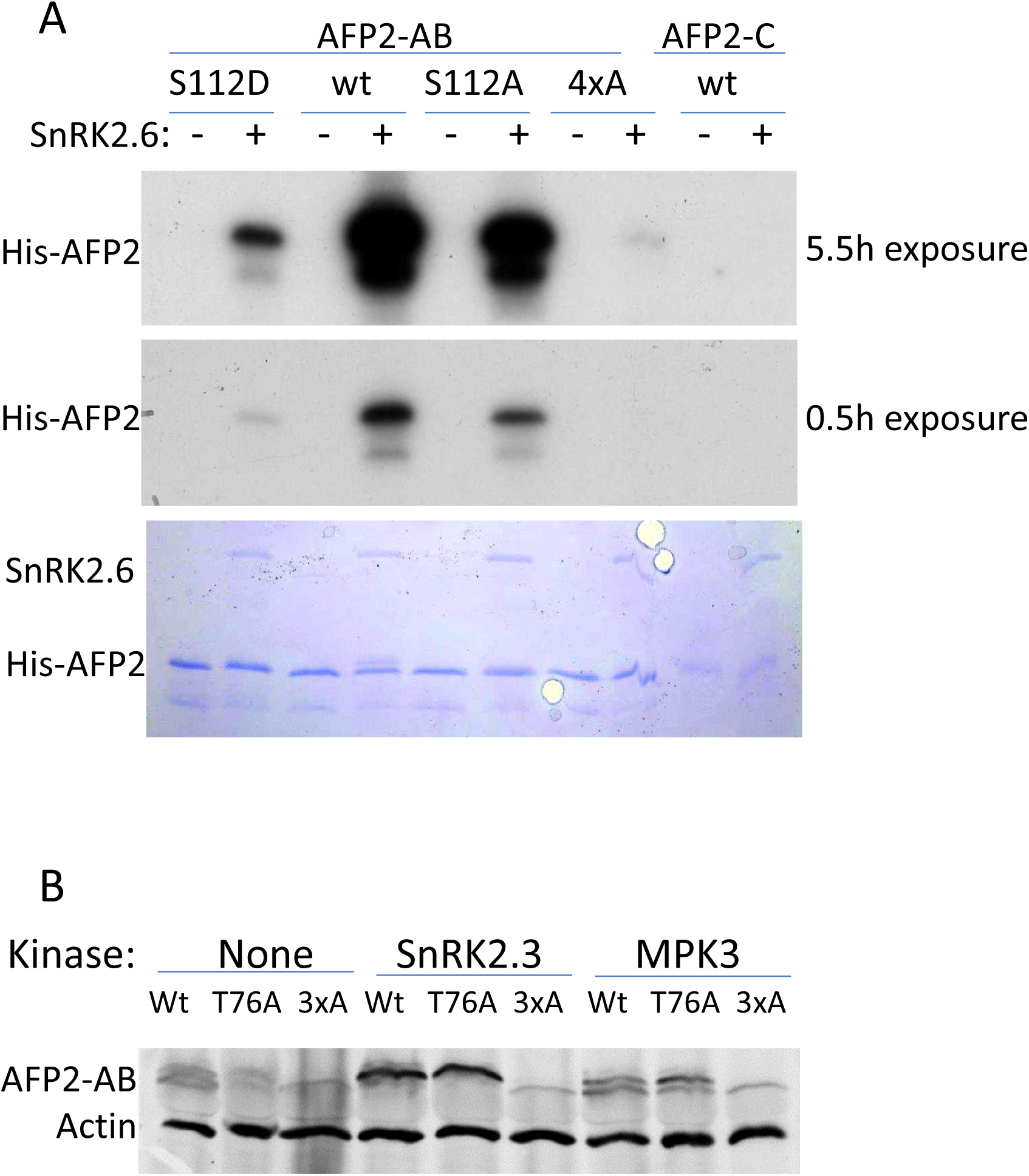
Tests of potential phosphorylation sites by kinase reactions with mutant AFP substrates. (A) *In vitro* kinase assay comparing phosphorylation of wild-type AFP2-AB domain with phosphomutants or AFP2-C domain. Top two panels are autoradiographs of the Coomassie-stained gel shown in the bottom panel. Affinity-purified GST-SnRK2.6 was pre-activated, then reactions set up as pool before aliquoting for combination with His-AFP2 substrates. Coomassie shows loading of AB fusions and SnRK. “4xA” = (S85,T111,S112,S129)A; as shown in Fig. 5E, (S85,S112)A gives the same result as the 4xA mutant. (B) Comparison of wild type and phosphomutant YFP-AFP2-AB domain mobility when transiently expressed in *N. benthamiana*, either alone or in combination with either Myc-SnRK2.3 or Myc-MPK3. “3xA” = (T76,S85,S112)A

**Supp. Fig. 12.**
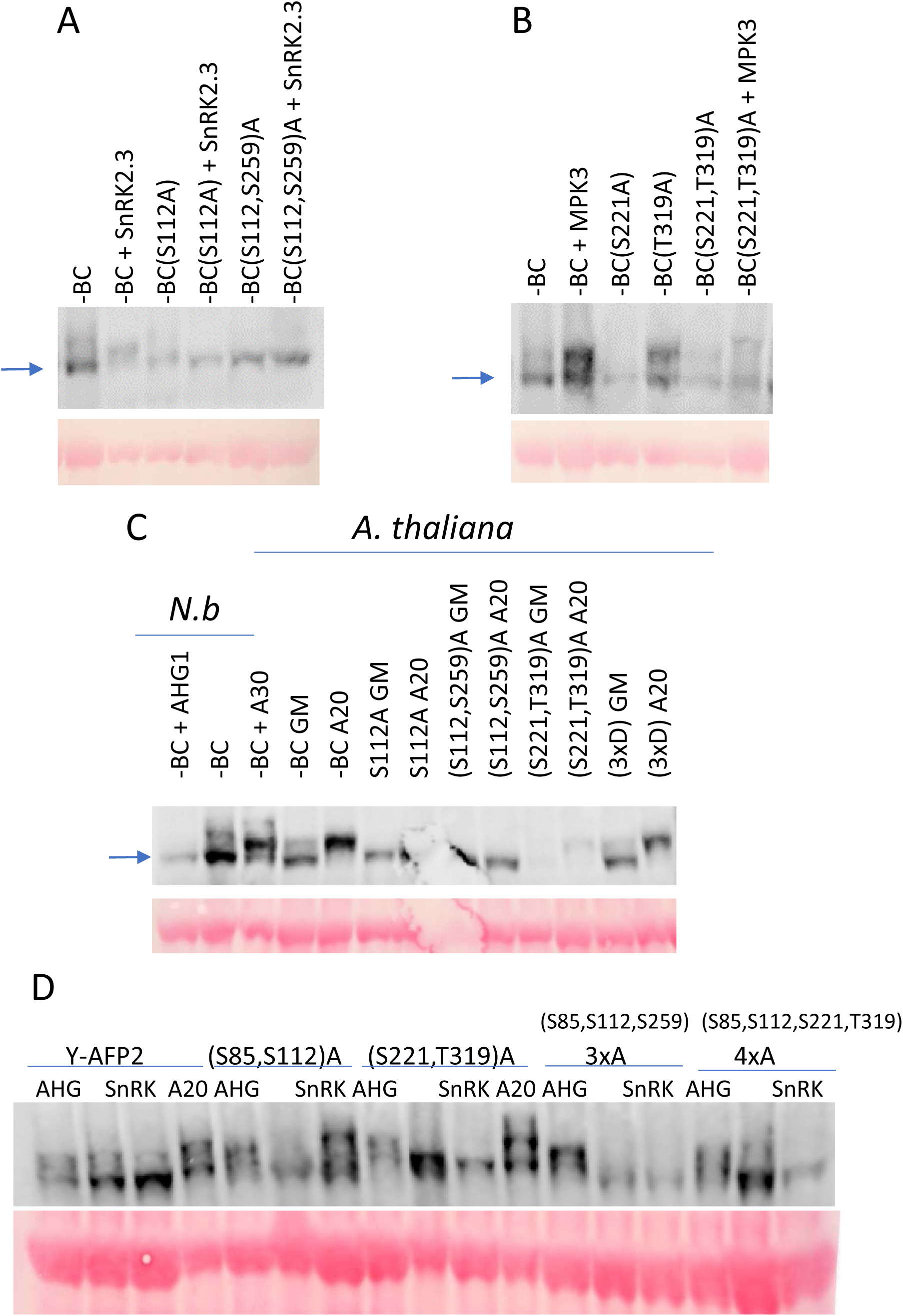
Comparison of phosphorylation status of AFP2-BC domain by Phos-tag gels. YFP-AFP2-BC fusions were expressed transiently in *N.benthamiana* (A,B,C) or stably in *A.thaliana* (C). Fusions were expressed alone or in combination with Myc-SnRK2.3 (A), Myc-MPK3 (B) or Myc-AHG1 (C). ABA or GM incubations were as described in Fig. 6AB. The “3xD” mutant is altered at S112, S221, and T319. The unphosphorylated AFP2-BC domain is identified by the arrows. Full-length AFP2 fusions were expressed transiently alone or in combination with Myc-AHG1 or Myc-SnRK2.3 in *N.benthamiana* (D).

**Supp. Fig. 13.**
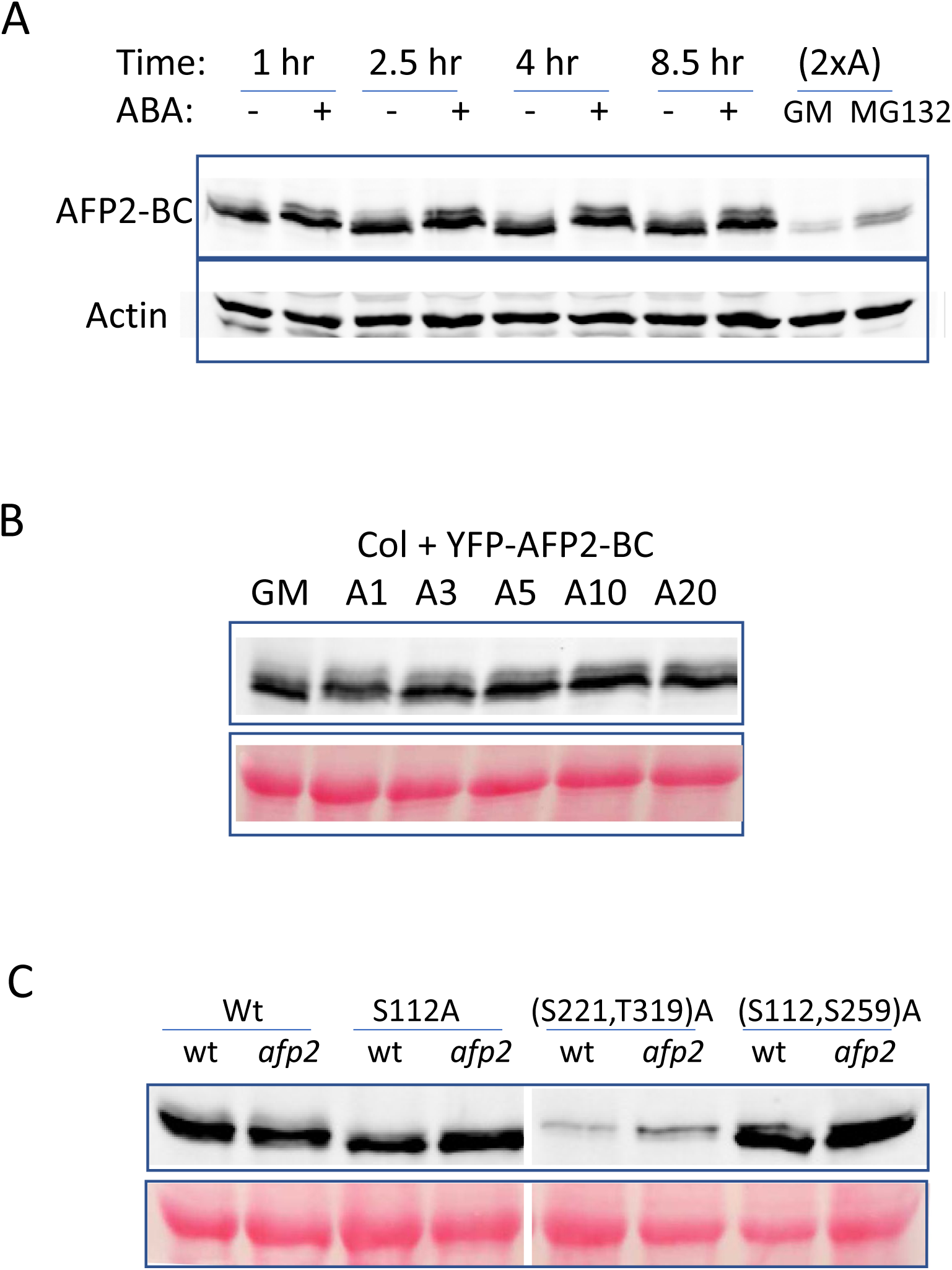
Effects of ABA, mutations and genetic background on expression and mobility of YFP-AFP2-BC transgenes in stably transformed *A. thaliana*. (A) Time course of changes in mobility of wild-type AFP2-BC in response to 20 μM ABA. (B) Dose response of ABA-induced changes in mobility of wild-type AFP2-BC domain in stably transformed *A. thaliana* incubated for 6 hrs in GM with or without ABA ranging from 1-20 μM. (C) Expression of wild-type (wt) YFP-AFP2-BC and transgenes with the indicated mutations in wt and *afp2* seedlings following germination and 7d growth on minimal media supplemented with 3 μM ABA. Upper panels are immunoblots probed with anti-GFP. Lower panels in B and C are the RbcL bands seen on Ponceau stained filters

**Supp. Fig. 14.**
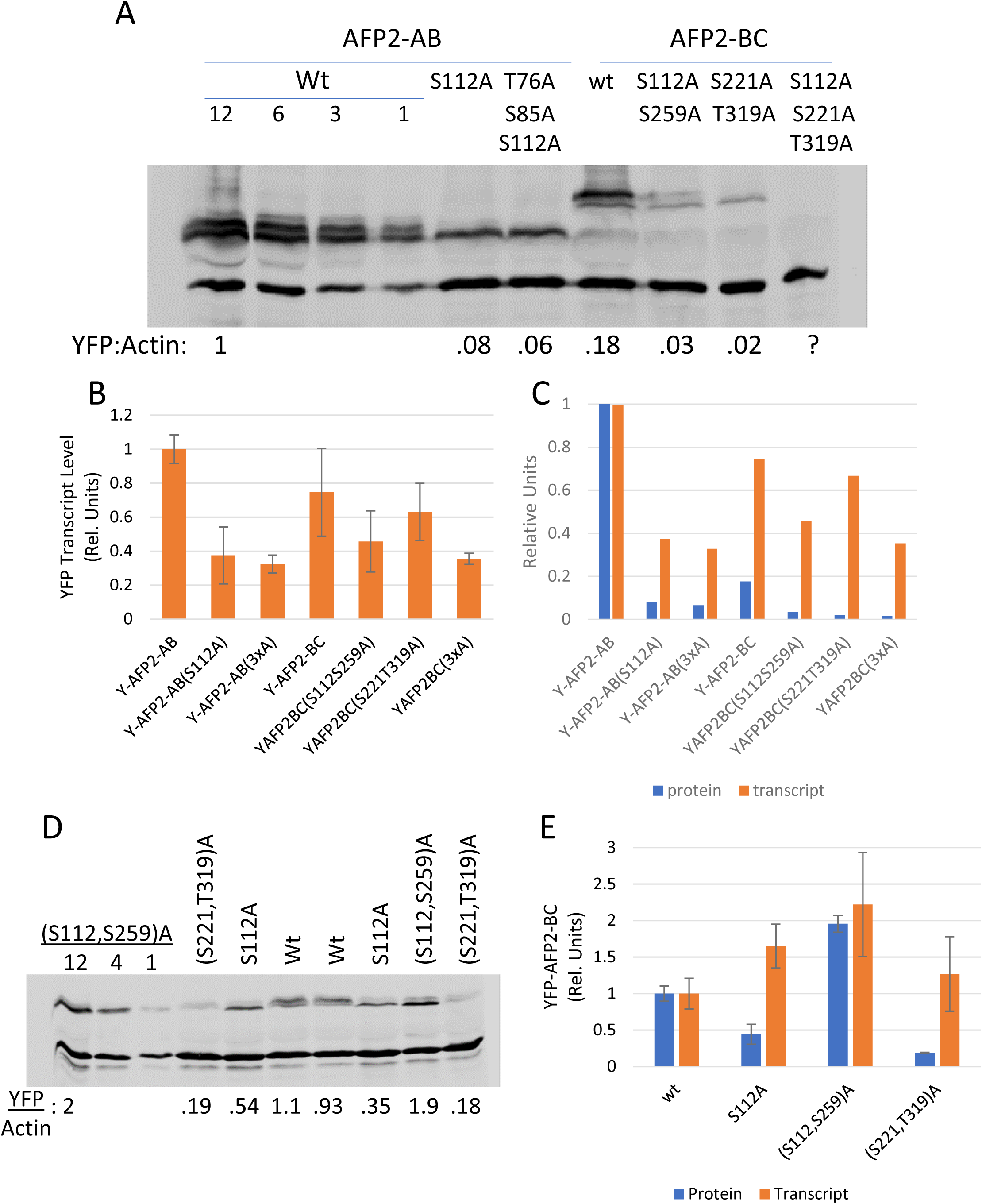
Comparison of protein and transcript levels for AFP2-AB or AFP2-BC domain fusions. Transient expression in *N. benthamiana* (A-C) or stable transformants in *A. thaliana* (D,E). The standard series of wt AFP2-AB shows the volume of extract loaded for the standard curve. Amounts of YFP fusion protein were normalized relative to actin. YFP-fusion transcripts were normalized relative to these stably expressed endogenous genes: PP2A (Niben101Scf09716g01002.1) of *N.benthamiana* and the geometric mean of PP2AA3 (At1g13320) and AP2M (At5g46630) of Arabidopsis.

**Supp. Fig. 15.**
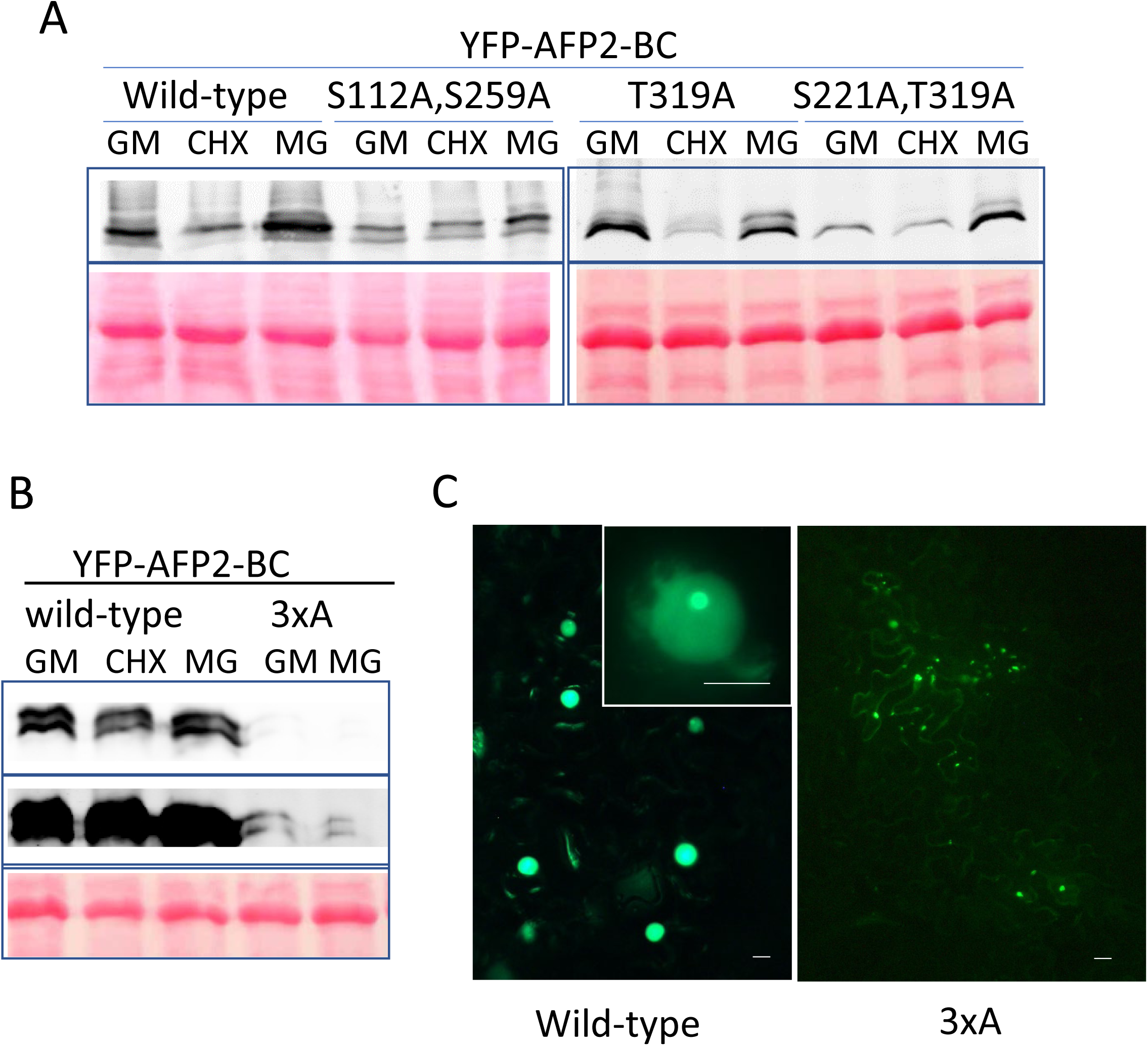
Comparison of stability and accumulation of transiently expressed wild type, double (S112A,S259A or S221A,T319A) or triple (3xA=S112A,S221A,T319A) mutant AFP2-BC domain fusions to YFP. Following infiltration, *N. benthamiana* leaf discs were infiltrated with GM (0.5x MS medium) with or without 100 μM cycloheximide (CHX) or MG132(MG), and incubated 6 hrs prior to harvest for extraction. Immunoblots were probed with anti-GFP (top panels) after staining with Ponceau (lower panels) (A,B). Micrographs of *N. benthamiana* leaves used for assay in B, taken with 10x objective on an Olympus AX70 epifluorescence microscope (C). Gain was set at 11.9 for 3xA mutant vs. 2.0 for wild-type. Inset is wild-type nucleus taken with 40x objective. Scale bars = 10 μm for wild-type, 20 μm for 3xA mutant

**Supp. Fig. 16.**
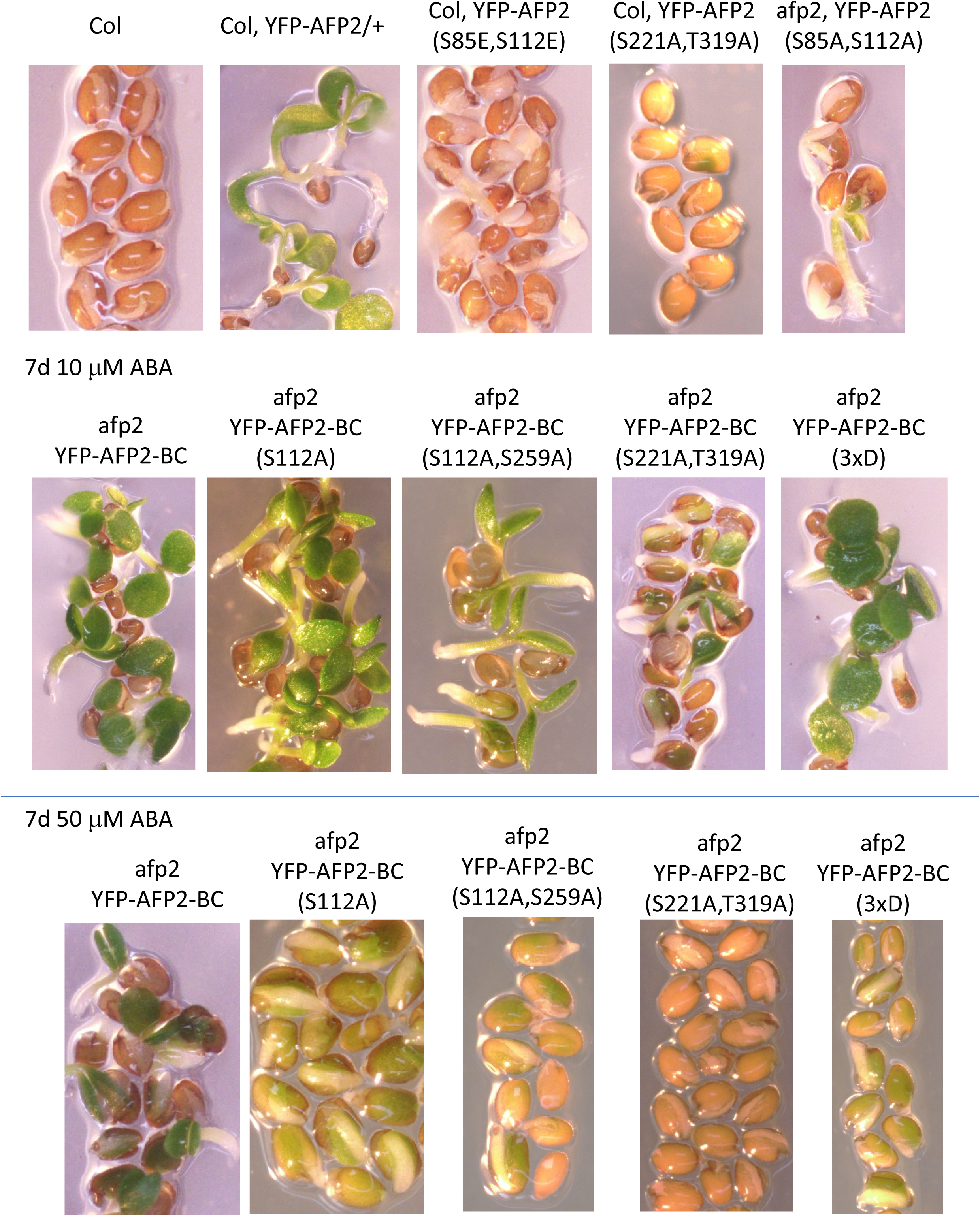
Germination phenotypes of wild type (Col) and transgenic seeds expressing the indicated YFP-AFP2 fusions after 7d incubation on either 10 or 50 μM ABA.

**Supp. Fig. 17.**
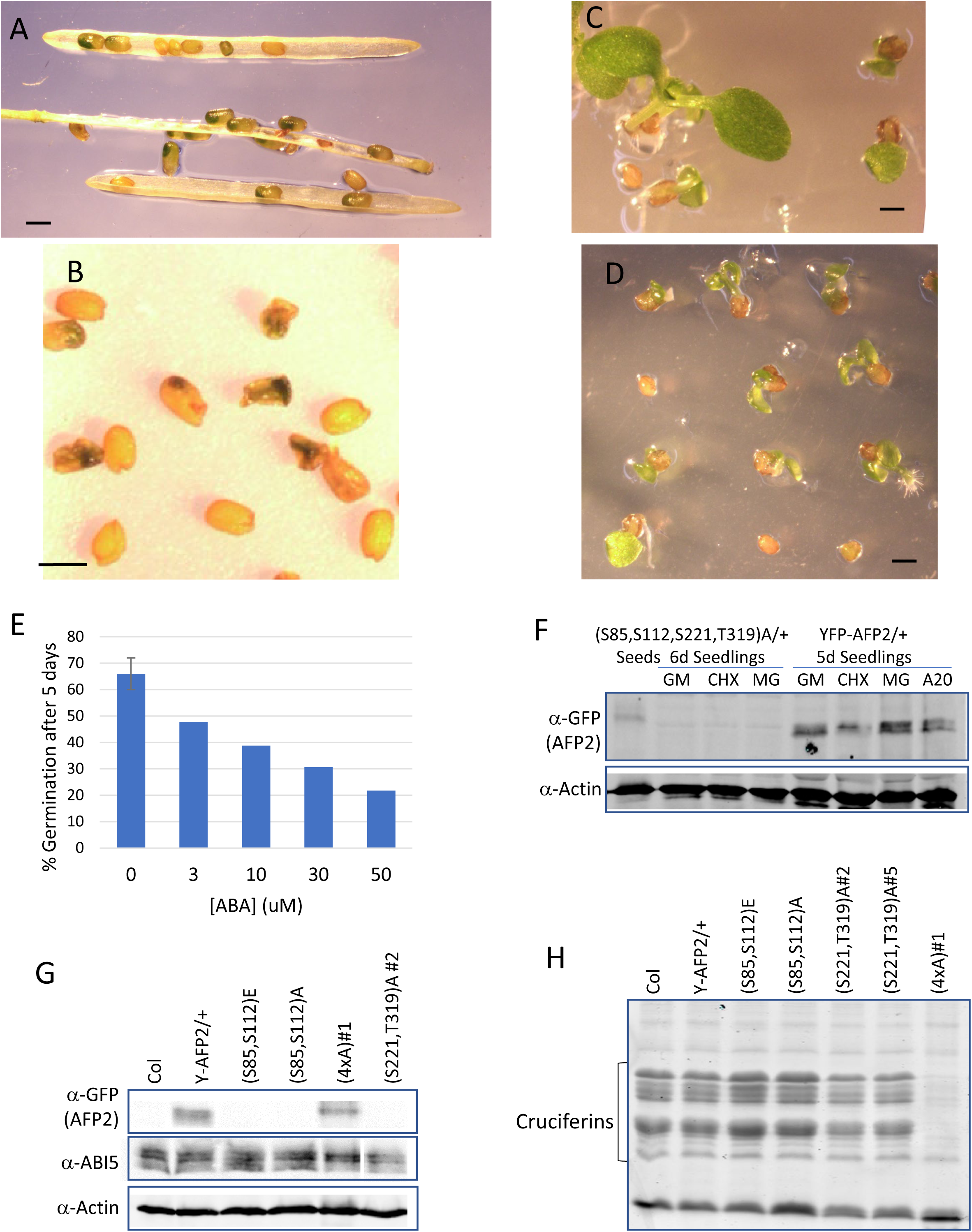
Transgenic line hemizygous for 35S-YFP-AFP2(S85,S112,S221,T319)A fails to complete seed maturation. (A) Maturing silique segregating green seeds. (B) Dry seeds with green embryos are desiccation intolerant and shrunken. (C) Comparison of seedlings from wild type siblings (left) and green seeds (right) removed from maturing siliques prior to desiccation. (D) Some green seeds fail to develop roots on minimal media. (E) Transgenic seeds are highly resistant to ABA-inhibition of germination. (F) Immunoblot comparing accumulation of wild type and mutant AFP2 fusions in seedlings following 6h incubation in GM with or without 100 μM CHX or MG132 or 20 μM ABA. Dry seed protein extracts of the indicated genotypes were separated on 10% SDS-PAGE for immunoblots (G) or 15% SDS-PAGE to resolve storage proteins (cruciferins) (H). “4xA” = 35S-YFP-AFP2(S85,S112,S221,T319)A. Scale bars = 0.5 mm

**Supp. Fig. 18.**
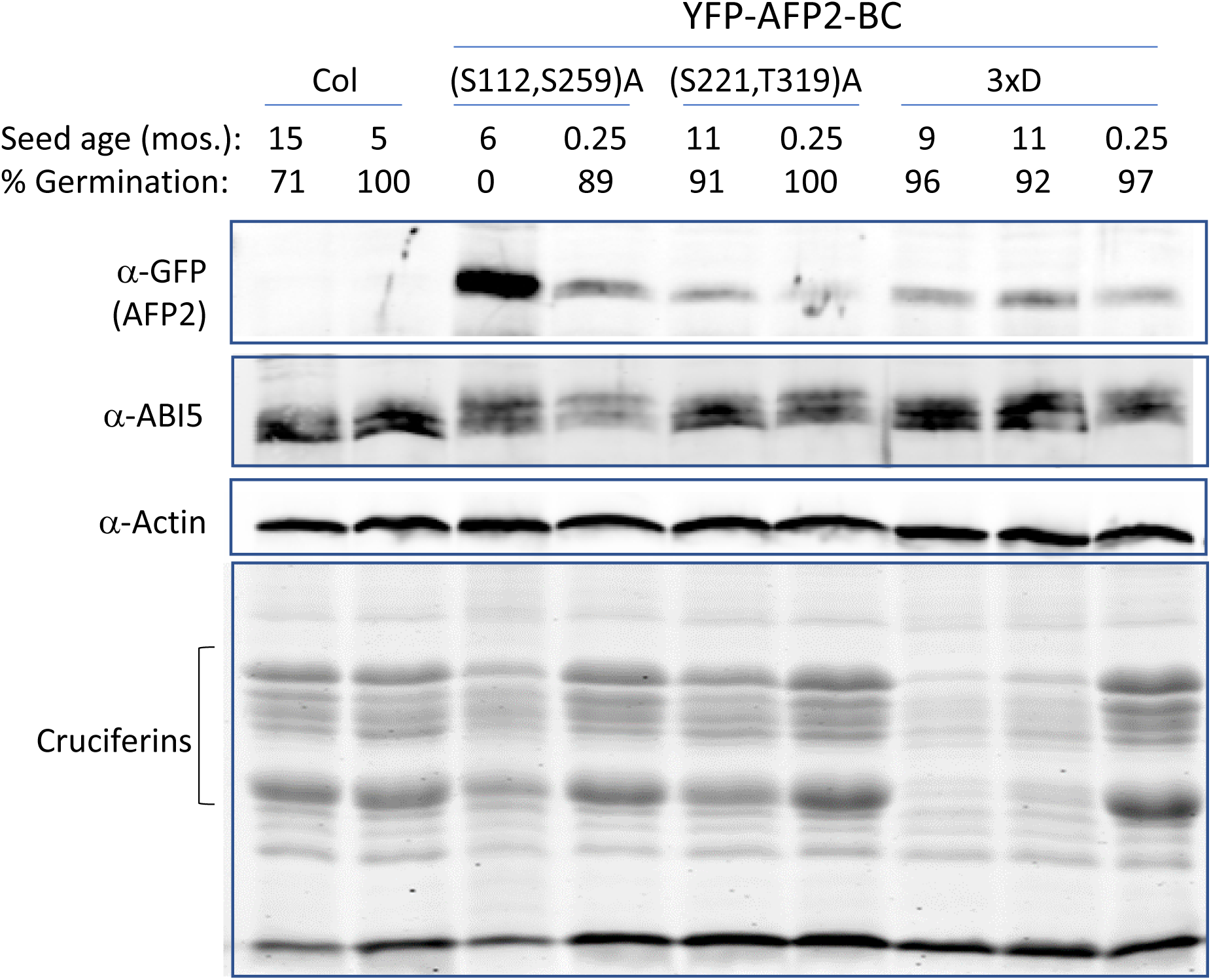
Effects of phosphomutant or phosphomimetic YFP-AFP2 fusions on storage protein accumulation and seed longevity. Dry seed protein extracts of the indicated genotypes and ages were separated on 10% SDS-PAGE for immunoblot or 15% SDS-PAGE for resolution of storage proteins. Germination was scored at 4d post-stratification incubation on minimal nutrient media.

**Supp. Fig. 19.**
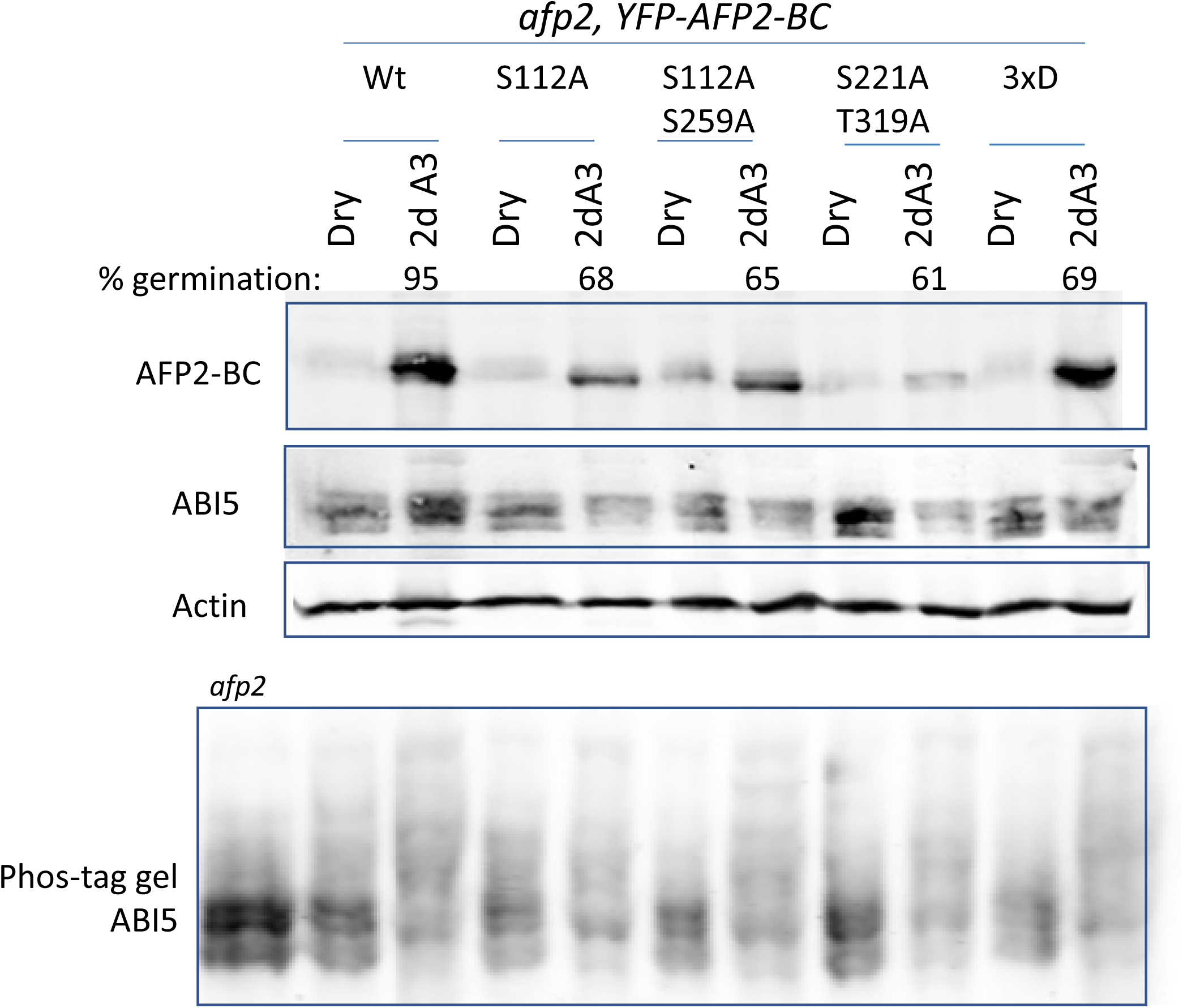
Effect of ABA on accumulation and phosphorylation of ABI5 in seeds with wild type or mutant YFP-AFP2-BC fusions in an *afp2* mutant background. Protein extracts were separated on 10% SDS-PAGE (upper panels) or a 7.5% Phos-tag gel, then transferred to filters for immunoblots. Germination on minimal media supplemented with 3 μM ABA (A3) was scored just prior to harvest.

**Supp. Fig. 20.**
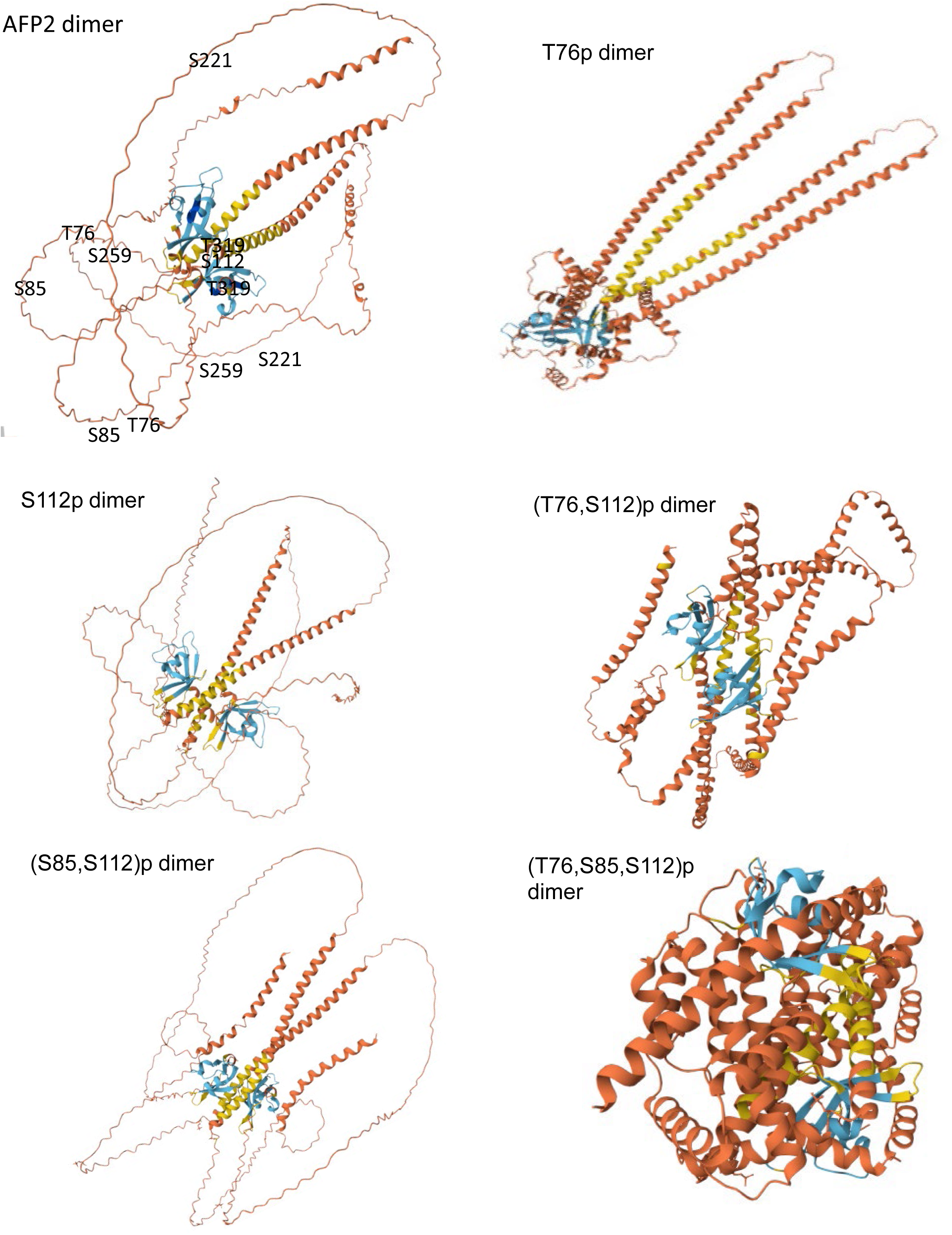
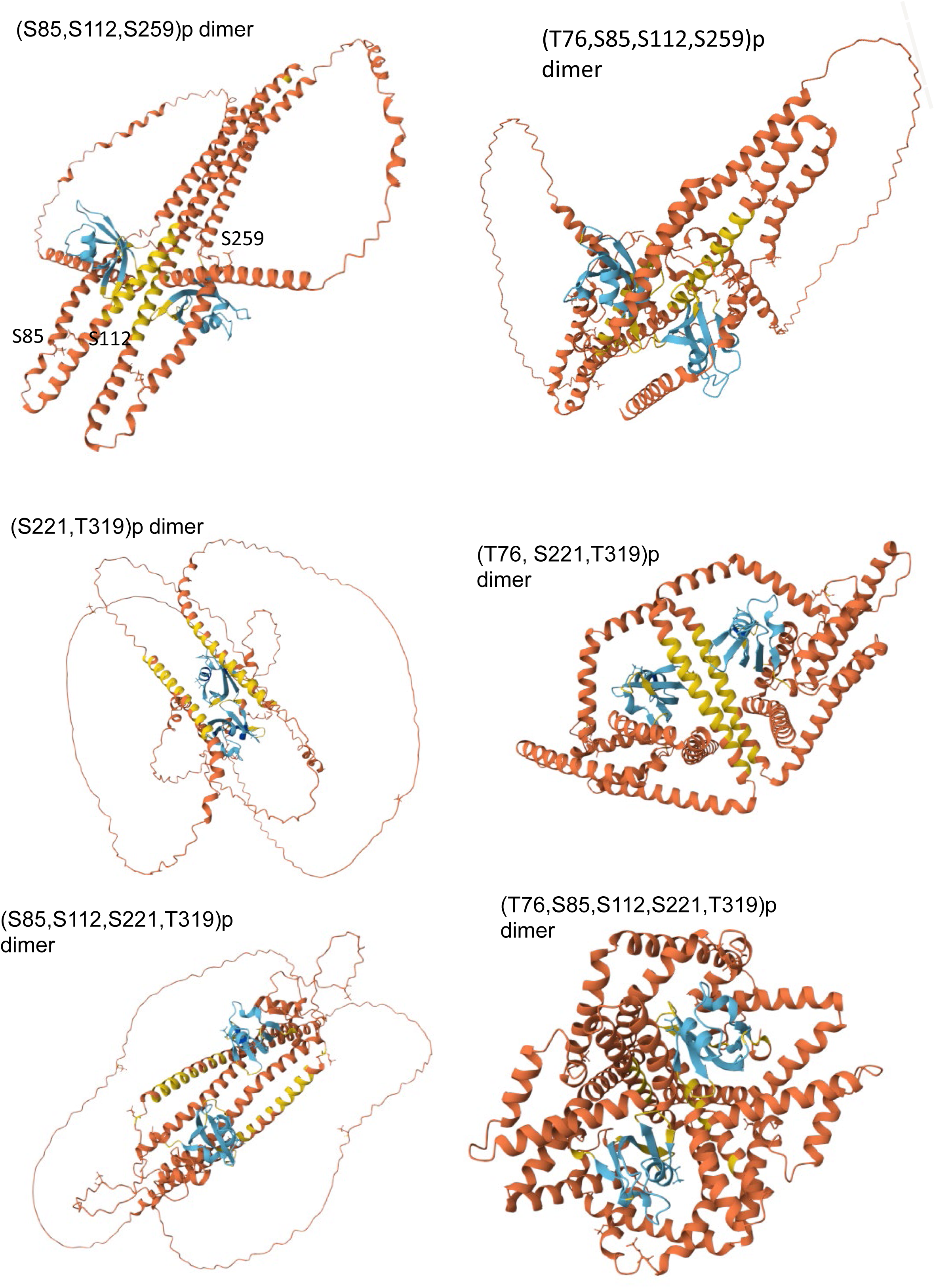
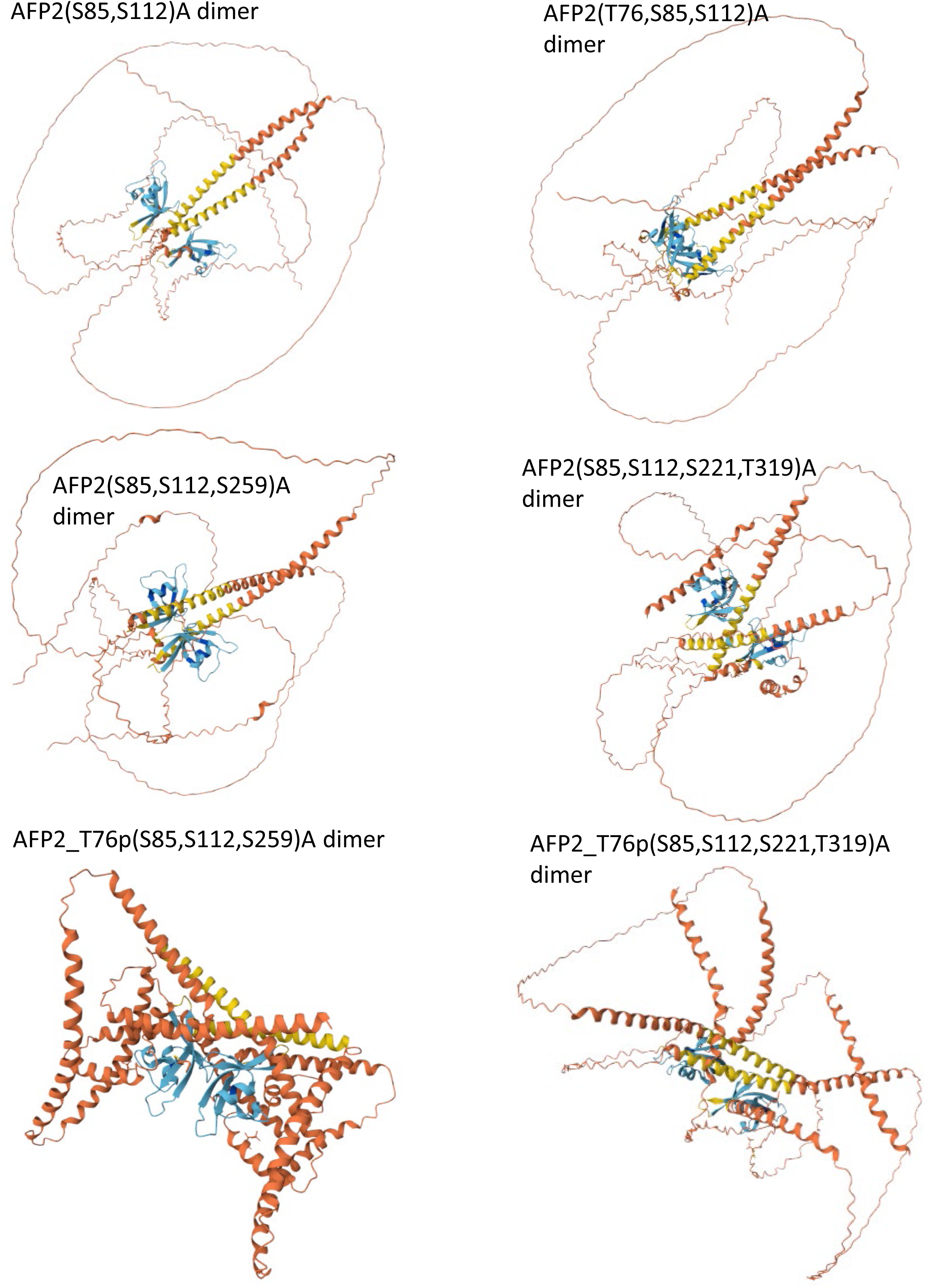
Alphafold predictions of AFP2 dimer structure when phosphorylated, or blocked from phosphorylation, at residues tested in this work.

## Notes

### Competing Interest Statement

The authors have declared no competing interest.

### Summary of Updates

Revision submitted to Plant Physiology on 7/25/2025.

